# MicroRNA-29 acutely regulates Memory Stability, Expression of Synaptic Genes, and DNA Methylation in the Mouse Adult Hippocampus

**DOI:** 10.1101/2024.11.12.623333

**Authors:** Aurelia Viglione, Chiara Giannuzzi, Elena Putignano, Raffaele Mario Mazziotti, Sara Bagnoli, Paola Tognini, Alessandro Cellerino, Tommaso Pizzorusso

## Abstract

MicroRNAs are key regulators of brain gene expression, with miR-29 family notably upregulated from development to adulthood and in aging, and showing links to cognitive decline. However, the extent to which miR-29 levels influence learning and memory processes, and its molecular mediators, remains to be determined. Here, we down- and up-regulated miR-29 levels in the dorsal hippocampus of adult mice to reveal miR-29 role in memory. Inhibition of miR-29 enhanced trace fear memory stability, increased Dnmt3a levels, and promoted DNA methylation in a DNMT3a-dependent manner. In contrast, increasing miR-29 impaired memory performances and decreased Dnmt3a levels, suggesting a destabilization of memory processes. Proteomic and transcriptomic analysis demonstrated that miR-29 antagonism upregulated RNA-binding and synaptic proteins and downregulated inflammation and myelin associated proteins. These results underscore miR-29’s pivotal role in memory persistence, plasticity, and cognitive aging, suggesting that miR-29 modulation could offer potential strategies for cognitive enhancement and age-related memory decline.

## Introduction

MicroRNAs (miRNAs) are small non-coding RNA molecules that modulate gene expression by binding to complementary sequences in target mRNAs, leading to the repression of translation and transcript destabilization [1].Through their ability to regulate hundreds to thousands of targets simultaneously, miRNAs enable fine-tuned, reversible control of gene expression programs. This regulatory flexibility is particularly important in the nervous system, where dynamic and coordinated changes in gene expression underlie synaptic plasticity, learning, and memory formation [2, 3].

One particular miRNA family, miR-29, has emerged as a potential regulator at the intersection of aging and memory. MiR-29 is extensively involved in a wide range of biological processes relevant to brain function such as apoptosis [4], metabolism [5], neuronal maturation [4], immunity [6, 7], and the aging-associated accumulation of iron in the brain [4, 8]. Notably, miR-29 expression increases across the lifespan in multiple tissues [9], including the nervous system of multiple species [10–12], and altered miR-29 levels have been associated with cognitive impairment [13]. However, the functional implications of these expression changes for neuronal plasticity and memory remain incompletely understood. Previous studies have reported seemingly opposing functional roles for miR-29 in the brain. Partial reduction of miR-29 levels has been shown to reduce lifespan, induce neurodegeneration, and promote molecular hallmarks associated with brain dysfunction [4, 8, 14]. On the other hand, however, overexpression of miR-29 has even been shown to induce premature aging in mice, suggesting that miR-29 is a driver of the aging process [14].

The miR-29 family, which includes miR-29a, miR-29b, and miR-29c, is encoded by two genomic loci: miR-29a/b-1 and miR-29b-2/c. Among these, miR-29a-3p (hereafter miR-29a) is particularly noteworthy, as it is the most upregulated during postnatal development and maturation of the mouse visual cortex [15] and its downregulation has been shown to slow cognitive decline and reduce beta-amyloid deposition [16]. At the molecular level, miR-29a could influence memory by modulating genes related to extracellular matrix and transcription regulation [15]. In particular, miR-29a directly targets DNA methyltransferases (DNMTs) [17–19], enzymes responsible for establishing and maintaining DNA methylation patterns. DNA methylation is a critical epigenetic mechanism for regulating gene expression in the brain and has been implicated in synaptic plasticity, learning, and memory [20, 21]. Rapid and dynamic changes in DNA methylation have been observed in response to neuronal activity [22, 23], and studies have demonstrated that DNA methylation writers are necessary for learning and memory stability [24–26]. Moreover, within the brain-expressed miRNAs, miR-29a emerges as notably synaptically enriched [27], suggesting the involvement of miR-29a in the regulatory network underpinning synaptic plasticity and the formation of memories [28]. However, the precise mechanisms through which miR-29a influences memory performance during development and in the adult brain remain elusive.

In particular, whether the age-dependent increase in miR-29 expression is protective or disruptive for cognitive function is still to be ascertained. Our study shows that inhibiting miR-29a using Locked Nucleic Acid (LNA) antagomirs (anti-miR29a) resulted in increased DNMT3a levels and improved memory retention, while enhancing miR-29a levels had the opposite effect. These effects could also be attributed to miR-29c, as anti-miR-29a may target both isoforms due to their sequences differing by only one nucleotide and sharing the same seed region, thereby largely regulating the same targets. These findings suggest that miR-29a not only plays a role in age-related cognitive decline but also exerts a bidirectional control over memory possibly by altering the epigenetic landscape. Indeed, we observe increased CpG methylation in promoter and exon regions associated with CpG islands and modulation of the hippocampal transcriptome and proteome. These effects of miR-29a antagonization on DNA methylation were blocked by down regulation of *Dnmt3a*. Our findings provide compelling evidence that miR-29a orchestrates the molecular changes necessary for the maintenance of emotional memories, highlighting its potential role in age-related cognitive decline.

## Results

### Age-Dependent increase of miR-29a levels correlates with *Dnmt3a* reduction in the dorsal hippocampus

To investigate the temporal expression pattern of miR-29a in the dorsal hippocampus, we conducted a quantitative PCR (qPCR) analysis at different time points. Our results revealed a significant 10-fold increase in miR-29a levels after postnatal day 10 (P10), reaching a plateau around P60 (Figure 1A). Because expression levels remain stable after this peak, we conducted the subsequent experiments at P90 to ensure that our analyses captured robust and sustained miR-29a expression. Interestingly, the specific temporal window of miR-29a increase aligns with critical periods characterized by heightened plasticity in various brain regions, accompanied by an increased sensitivity of chromatin configuration to environmental stimuli [29, 30]. Simultaneously, we examined the age-related modulation of *Dnmt3a*, a well-validated target of miR-29 [18, 31]. With increasing age, *Dnmt3a* expression showed a significant downregulation (Figure 1B), displaying a subject-by-subject negative correlation with miR-29a levels (Figure 1C). In a previous study, a comparable expression pattern was reported for the visual cortex, wherein the age-related elevation of miR-29a and decrease in *Dnmt3a* expression levels were associated with a significant decrease in ocular dominance plasticity and the expression of molecular critical period regulators [15]. Given that age-related decline in *Dnmt3a* levels has been proposed as a potential mechanism contributing to age-related cognitive impairments [32], we decided to explore the involvement of miR-29a in modulating fear memories. To evaluate whether and how the reduction in miR-29a expression affects fear memory formation and extinction, we injected a LNA oligonucleotide, whose sequence is complementary to miR-29a (anti-miR29a), or scrambled oligonucleotide (scr) in the dorsal hippocampus of adult mice, and then we tested hippocampal memories through a hippocampus dependent trace fear conditioning protocol (TFC) (Figure 1D). As previously observed using LNA oligonucleotides against miR-29a in the visual cortex [15], anti-miR29a demonstrated a stable and specific binding affinity to their complementary miRNA, effectively blocking its activity, and exhibiting resistance to enzymatic degradation. These attributes render LNAs highly tolerable and validated, as underscored by their approval for clinical use in human subjects [33–35]. As expected, the qPCR revealed that anti-miR29a treatment caused a decrease in miR-29a (Figure 1E) and an increase in the *Dnmt3a* (Figure 1F). We also found a sample-by-sample negative correlation between *Dnmt3a* and miR-29a levels after LNA or scr injection (Supplementary Figure 1A) confirming the tight regulation of *Dnmt3a* levels by miR-29a. To control for the spatial spreading of our treatment, we quantified *Dnmt3a* levels also in the ventral hippocampus. The results revealed no significant difference in *Dnmt3a* expression (Supplementary Figure 1B), confirming the specificity of the injection effects within the dorsal hippocampus. No effect was observed on other factors related to DNA methylation such as MeCP2 and Tet3 (Supplementary Figure 2A-D). To assess specificity of the treatment among miR-29 isoforms, we assessed the expression of miR-29b and miR-29c in anti-miR29a treated animals. We found that miR-29c expression was strongly reduced, while no significant alterations were detected in miR-29b expression (Supplementary Figure 3A-B). This result is consistent with the fact that miR-29a and miR-29c share an identical seed sequence and differ by only one nucleotide located outside of the seed region and regulate, regulating largely overlapping target transcripts.

**Figure 1:**
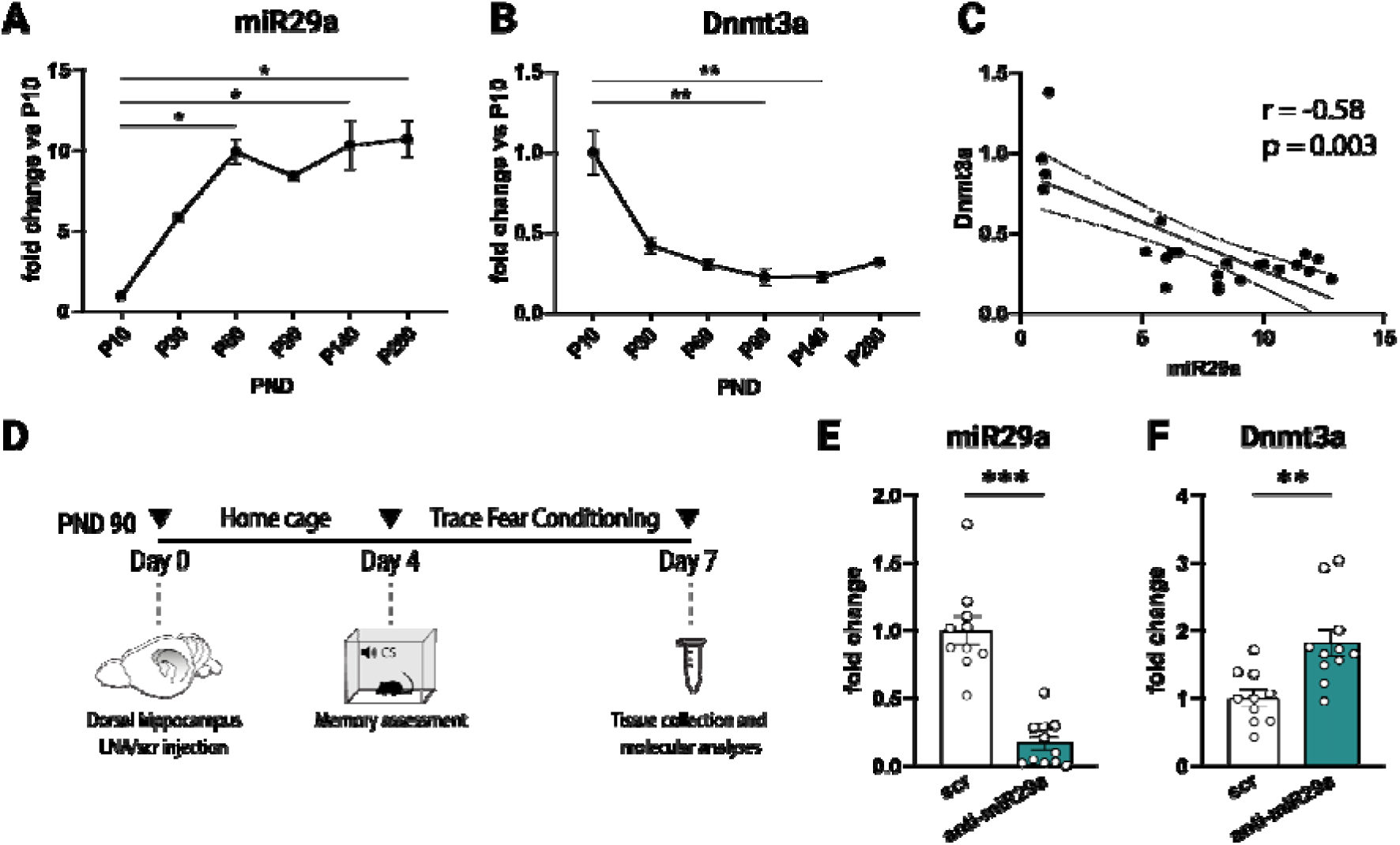
Age-dependent increase of miR-29a correlates with *Dnmt3a* expression in the dorsal hippocampus. **(A)** Dorsal hippocampus miR-29a expression level (normalized to P10) at different ages (Kruskal-Wallis: H(5) = 16.22, P *<* 0.006; Dunn’s multiple comparison vs P10: *P *<* 0.05 , n = 3/4 mice for each age). **(B)** Dorsal hippocampus *Dnmt3a* expression level (normalized to P10) at different ages (Kruskal-Wallis: H(5) = 17.10, P *<* 0.004; Dunn’s multiple comparison vs P10: **P *<* 0.01; n = 3/4 mice for each age). **(C)** Animal by animal correlation between normalized miR-29a and *Dnmt3a* levels in the dorsal hippocampus at different ages (Spearman’s correlation). **(D)** Experimental design for LNA anti-miR29a treatment in the dorsal hippocampus. **(E)** Effects of anti-miR29a treatment on miR-29a expression. Fold change values normalized to scr treated animals (scr: N = 10, anti-miR29a N = 11; Mann-Whitney U test: U = 1, p *<* 0.0001). **(F)** Effects of LNA-anti-miR29a treatment on *Dnmt3a* expression. Fold change values normalized to scr treated animals (scr: N = 10, anti-miR29a N = 11; Mann-Whitney test: U = 13, p *<* 0.01). PND = Postnatal Day.

### MiR-29 downregulation promotes memory stability during fear extinction

Three days after anti-miR29a injection, we conditioned animals using a TFC protocol. During TFC the neutral conditioned stimulus (CS, tone) and the aversive unconditioned stimulus (US, shock) are separated in time by a trace interval (trace). The absence of contiguity between the tone and the shock critically involves the hippocampus and this protocol can be used to evaluate hippocampal dependent learning and memory [36] (Figure 2A). We observed that the treatment did not affect learning acquisition, since both anti-miR29a and scr treated mice showed similar freezing during learning (Figure 2B and Supplementary Figure 4A-C). When testing freezing responses 24h after training, the two groups still did not differ in the percentage of time freezing (Figure 2D). However, on the first day of the extinction protocol (day 3, Early Extinction) we observed that anti-miR29a treated mice showed a higher fear retention, as compared to the control group (Figure 2E). This observation suggests that miR-29a developmental upregulation affects strength and persistence of memory, but it does not affect memory formation.

**Figure 2:**
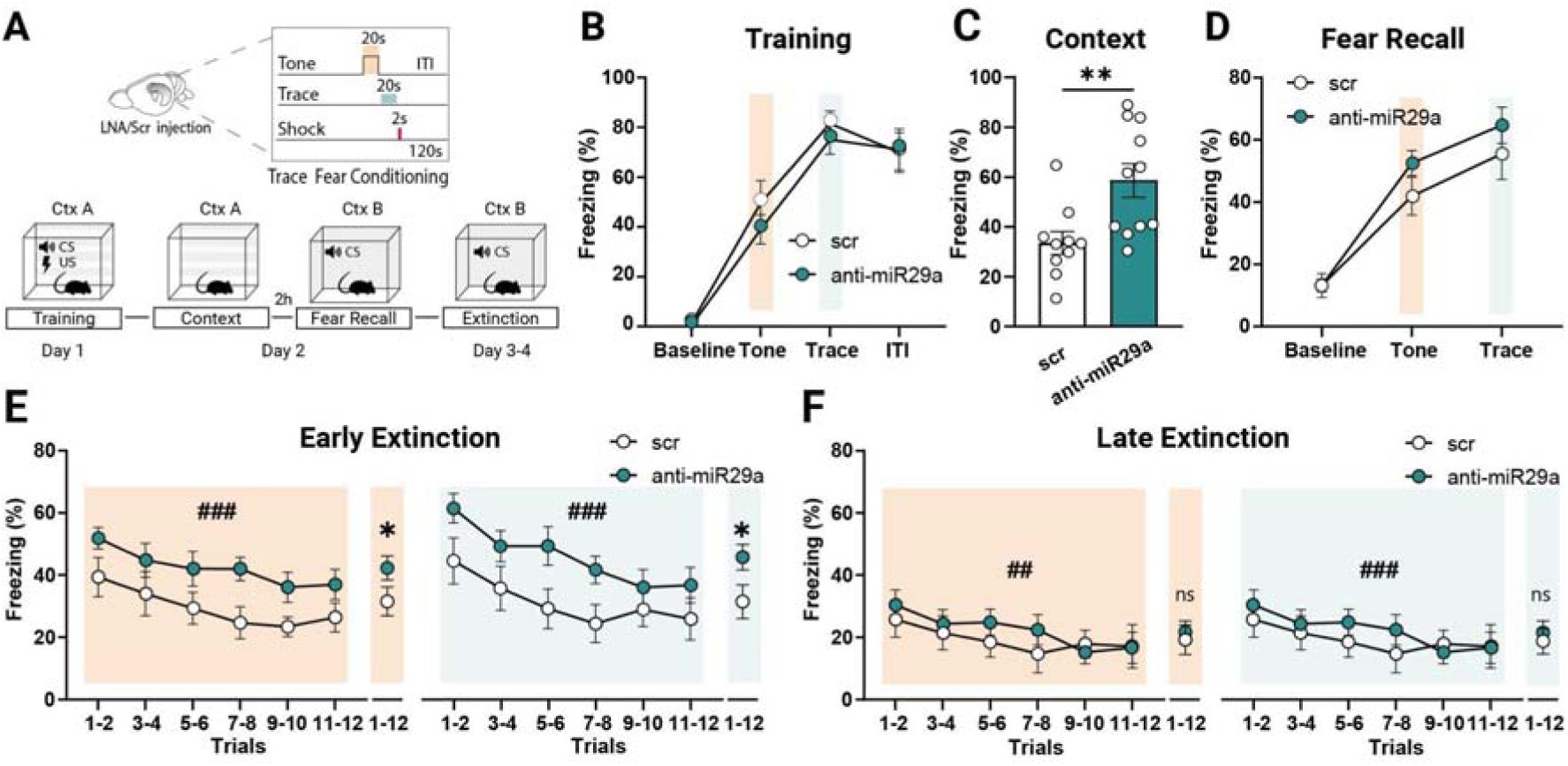
Reducing hippocampal miR-29a expression promotes memory stability. **(A)** Diagram showing the TFC paradigm and timeline. **(B)** Anti-miR29a and scr-treated animals showed no differences at baseline or in fear memory acquisition, as indicated by similar freezing responses during the final two-tone, trace trials and ITI of the TFC training session (Two-way ANOVA, test interval × treatment interaction F(3, 76) = 0.39, p = 0.76; main effect of test interval F(3, 76) = 64.33, p *<* 0.0001; main effect of treatment, F(1, 76) = 0.93, p = 0.34). **(C)** Downregulation of miR-29a in the dorsal hippocampus enhances contextual memory recall (Mann-Whitney U test: U = 16, p *<* 0.01, scr: N = 10, anti-miR29a N = 11). **(D)** No treatment differences were observed at baseline or during the final two-tone and trace trials of the TFC fear recall session (Two-way ANOVA, test interval × treatment interaction F(2, 57) = 0.59, p = 0.56; main effect of test interval F(2, 57) = 41.74, p *<* 0.0001; main effect of treatment, F(1, 57) = 2.33, p = 0.13). **(E)** Percentage of freezing during Early extinction. anti-miR29a mice showed a higher fear retention compared to the control group during both tone (Two-way RM ANOVA, main effect of treatment F(1, 19) = 4.78, p *<* 0.05; main effect of trials F (2.98, 56.58) = 6.69, p *<* 0.0001; trials × treatment interaction F(5, 95) = 0.29, p = 0.92) and trace intervals (Two-way RM ANOVA, main effect of treatment F(1, 19) = 4.47, p *<* 0.05; main effect of trials F(3.26, 62.02) = 9.36, p *<* 0.0001; trials × treatment interaction F(5, 95) = 0.77, p = 0.57). **(F)** During the Late extinction, anti-miR29a treated mice showed no significant differences in the percentage of freezing compared with controls (Tone: Two-way RM ANOVA, main effect of treatment F(1, 18) = 0.19, p = 0.66; main effect of trials F (2.56, 46.16) = 4.81, p *<* 0.01; trials × treatment interaction F(5, 90) = 0.87, p = 0.50; Trace: Two-way RM ANOVA, main effect of treatment F(1, 18) = 0.24, p = 0.63; main effect of trials F(3.95, 71.15) = 6.26, p *<* 0.001; trials × treatment interaction F(5, 90) = 2.12, p = 0.07). scr: N = 10, anti-miR29a N = 11; *indicates the main effect of treatment, #indicates the main effect of trials. CS= conditioned stimulus

We also assessed freezing behavior induced by exposure to the conditioned context (Figure 2A). Remarkably, mice subjected to anti-miR29a treatment exhibited a pronounced enhancement in their response to the contextual cues compared with the scr control group (Figure 2C). Despite this general amplification in fear response, anti-miR29a treated mice exhibited a typical extinction profile. Following the second extinction session (Late Extinction), both anti-miR29a and scr treated mice demonstrated significantly reduced freezing responses (Figure 2F). These findings offer insights into miR-29 role in modulating the persistence of fear memories. This effect may be mediated through the regulation of miR-29 target genes, particularly those involved in DNA methylation pathways and gene expression control.

### MiR-29 inhibition alters DNA methylation patterns and influences memory-associated pathways at the transcript and protein level

We employed reduced representation bisulfite sequencing (RRBS) to investigate changes in DNA methylation patterns induced by anti-miR29a treatment in comparison to scr treated hippocampal samples. RRBS allows for an exploration of methylation alterations in a CpG-enriched subset of the genome, primarily focusing on promoters [22, 37]. We analyzed differential methylation of the sequenced CpGs and selected differentially methylated CpGs (DMCs, q<0.05) with a difference greater than 5% in absolute value. Our analysis identified 35,246 DMCs after anti-miR29a treatment, with a notable prevalence of hypermethylation (32,878 CpGs) over hypomethylation (2,368 CpGs) (Figure 3A). To test whether these methylation changes are mediated by DNMT3a, we analyzed hippocampal samples treated with anti-*Dnmt3a* alone or in combination with anti-miR29a. Anti-*Dnmt3a* treatment resulted in 1,742 hypermethylated and 2,082 hypomethylated CpGs, showing a bias toward hypomethylation. The combined treatment produced 3,491 hypermethylated and 6,143 hypomethylated CpGs, displaying a similar hypomethylation-skewed profile (Figure 3A). These results indicate that DNMT3a inhibition counteracts the hypermethylation bias induced by anti-miR29a and suggest that a substantial fraction of miR-29a-dependent methylation changes are mediated through DNMT3a. Partial least squares discriminant analysis (PLS–DA) of DNA methylation profiles revealed clear separation of anti-miR29a samples from controls as well as from all other groups. Anti-*Dnmt3a* and combined treatment samples largely overlapped with each other but remained distinct from controls, which formed a separate cluster (Figure 3B). These findings suggest that DNMT3a activity contributes substantially to the methylation landscape induced by miR-29a inhibition. Comparison of DMCpGs between anti-miR29a alone and combined treatment revealed a highly non-random distribution of concordant and discordant CpGs (χ² = 473.11, p < 2.2 × 10L¹L). DMCpGs were classified based on their methylation changes in the two treatments into concordant and discordant quadrants. Concordant hypermethylated CpGs, which were hypermethylated in both anti-miR29a and combined treatment, were enriched for synaptic processes, including vesicle-mediated transport in synapse (Figure 3C). In contrast, discordant CpGs, which were hypermethylated in anti-miR29a but hypomethylated in the combined treatment, were the majority and were enriched for regulation of membrane potential, positive regulation of nervous system development, forebrain development, and cell junction assembly (Figure 3C). These results indicate that miR-29 inhibition affects multiple neuronal pathways: some epigenetic changes persist even when DNMT3a is inhibited, while others are reversed, highlighting a partial DNMT3a dependence of miR-29-mediated DNA methylation in hippocampal neurons.

**Figure 3:**
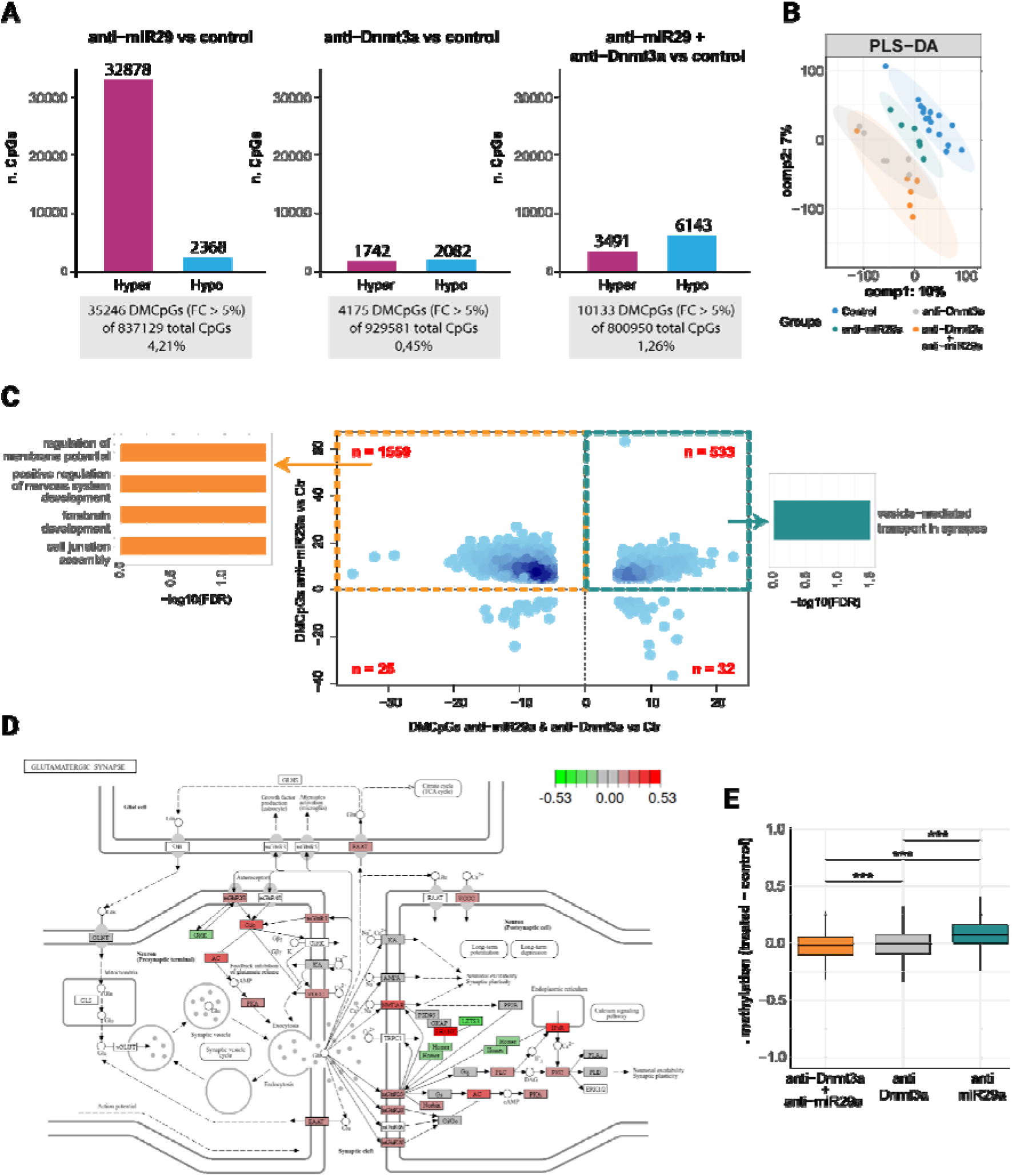
Dnmt3a mediates the epigenomic effects of miR-29a in the mouse dorsal hippocampus. **(A)** Differentially methylated CpGs (DMCpGs). Bar plots showing the number of significantly hypermethylated (magenta) and hypomethylated (blue) CpGs identified in each treatment compared to control (q ≤ 0.05; |Δmethylation| > 5%). Numbers above the bars indicate the absolute number of DMCpGs in each category. Grey boxes report the total number and percentage of DMCpGs relative to the total CpGs analyzed in each comparison. **(B)** Sample clustering based on DNA methylation profiles. PLS–DA analysis of genome-wide DNA methylation across experimental groups. Each dot represents one animal. The first two components are shown (comp1: 10% variance explained; comp2: 7%). Ellipses indicate the 95% confidence interval for each group, highlighting separation among treatments. **(C)** Differentially methylated CpGs and GO enrichment analysis across treatment conditions. Central panel: Scatter plot of DMCpGs comparing methylation changes in the combined treatment (anti-miR29a + anti-*Dnmt3a* vs control; x-axis) with those in anti-miR29a alone (vs control; y-axis). Each dot represents one DMCpG. Dashed lines indicate no change relative to control and define four quadrants. Quadrants 1 and 3 represent concordant CpGs (same direction of change), whereas quadrants 2 and 4 represent discordant CpGs (opposite direction). The number of CpGs per quadrant is shown in red. A χ² test revealed a significant enrichment of concordant CpGs compared to a uniform distribution (χ² = 473.11, p < 2.2e-16). Left and right panels: GO Biological Process enrichment analyses performed on CpGs from quadrant 2 and quadrant 1, respectively. Bars show significantly enriched GO terms, plotted as −log10(FDR). **(D)** Pathway mapping of differentially methylated genes. Schematic representation of the glutamatergic synapse pathway with genes associated with DMCpGs overlaid onto the pathway map. Nodes are color-coded according to the direction and magnitude of methylation change (green: hypomethylation; red: hypermethylation; color scale shown above). This analysis highlights coordinated methylation changes affecting multiple components of synaptic signaling and plasticity pathways. **(E)** Global CH methylation changes. Boxplots showing the distribution of Δ methylation (treated − control) at CH sites for each treatment group. Values represent per-animal differences between treated and contralateral control hemispheres. Statistical comparisons were performed using Welch’s ANOVA followed by Bonferroni-corrected pairwise t-tests. Horizontal lines indicate medians; boxes represent the interquartile range (IQR); whiskers extend to 1.5× IQR. All pairwise comparisons were significant (p < 0.001).

Gene Ontology (GO) enrichment analysis of hypermethylated CpGs identified after anti-miR29a treatment revealed enrichment for pathways related to synaptic signalling, extracellular matrix organisation, and kinase activity. Mapping methylation changes onto the KEGG glutamatergic synapse pathway (mmu04724) using Pathview showed widespread hypermethylation across multiple synaptic genes, suggesting that miR-29 regulates epigenetic programs involved in hippocampal-dependent plasticity and memory persistence (Figure 3D). Finally, analysis of global CH methylation differences across treatment groups revealed significant alterations, with all pairwise comparisons reaching statistical significance (Figure 3E), further supporting a DNMT3a contribution to the broader epigenomic effects induced by miR-29a inhibition. The enhanced DNA methylation in response to anti-miR29a treatment was confirmed also in an independent cohort of mice (Supplementary Figure 5).

To investigate the possible downstream consequences of epigenetic changes we performed RNAseq and mass-spectrometry-based proteomics (Figure 4). The analysis of differentially expressed proteins retrieved 420 significant protein groups out of 4878 detected protein groups. RNAseq detected 946 differentially expressed genes (DEGs). The analysis of the most overrepresented miRNA binding site in up-regulated DEGs revealed the expected enrichment for miR29 binding sites (miTEA [40], FDR < 10^-16^), confirming effective antagonism of miR-29 family. We first sought to identify pathways that regulated in a congruent direction both at the transcript and protein level, among these, glutamatergic transmission (KEGG mmu04724) stood out as glutamate receptors and transporters were upregulated often both at transcript and protein levels (Figure 4A). This pattern is clearly consistent with enhancement of synaptic plasticity, which is further supported by a large fraction of overrepresented GO terms related to synaptic function and plasticity, as well as neural development, among up-regulated genes (Supplementary Figure 6A). The analysis of downregulated genes revealed the strongest enrichment for genes expressed in microglia, followed by cells of both the myeloid and lymphoid lineage (Figure 4B). In addition, a striking down-regulation of genes coding for lysosomal protein (KEGG mmu04142) was observed (Figure 4C). This is further supported by a large fraction of overrepresented GO terms related to both adaptive and innate immunity among up-regulated genes (Supplementary Figure 6B). Downregulated proteins, on the other hand, showed a significant enrichment for genes expressed in oligodendrocytes only (Figure 4D) and coding for myelin sheath (GO 0043209).

**Figure 4.**
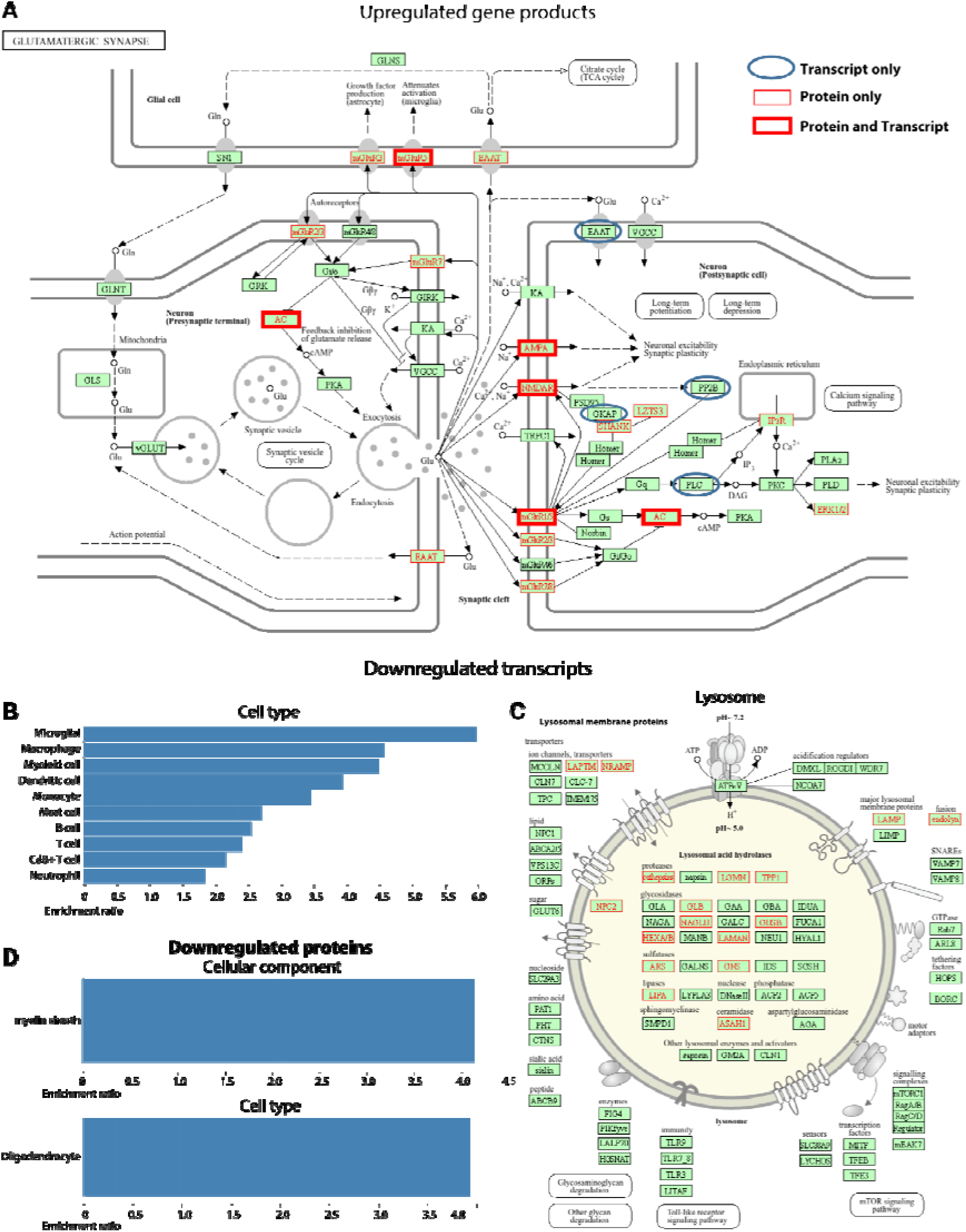
Integrated transcriptomic and proteomic analysis reveals synaptic potentiation and downregulation of immune and lysosomal programs. **(A)** Mapping of differentially expressed genes and proteins onto the KEGG glutamatergic synapse pathway (mmu04724). Ionotropic (e.g., NMDA, AMPA subunits), metabotropic receptors, and glutamate transporters are predominantly upregulated at the transcript and/or protein level, indicating enhanced synaptic plasticity. Colors indicate: transcript (blue), protein (red), both (solid red). **(B)** Cell-type enrichment analysis of downregulated genes shows strong overrepresentation of microglia-expressed genes, followed by myeloid and lymphoid cells. **(C)** Mapping of downregulated genes and proteins onto the KEGG lysosome pathway (mmu04142) reveals marked reduction of lysosomal hydrolases, membrane proteins, and acidification regulators. **(D)** Cell-type enrichment analysis of downregulated proteins highlights significant enrichment for oligodendrocyte proteins and myelin sheath components (GO:0043209).

To complement the analysis of GO term overrepresentation, we explored the interactome of regulated proteins and transcripts by analyzing the physical interactions of the upregulated differentially-expressed proteins (DEPs, FDR <0.1, Log2(fold change) > 0.1). We applied STRING network analysis [41] and found DEPs to be strongly connected (Supplementary Figure 7, expected number of edges: 295; number of edges: 938; PPI enrichment p-value:< 1.0e^-16^). The protein network consisted almost entirely of two tightly clustered communities of functionally-interacting proteins: one made of synaptic proteins and the second of RNA binding proteins. STRING network analysis found DEGs (FDR <0.1, DESeq2) to be strongly connected (Supplementary Figure 7, expected number of edges: 345; number of edges: 660; PPI enrichment p-value:< 1.0e-16). Similarly to DEP network, also the DEG network was strongly enriched for synaptic terms and RNA binding activity, indicating that the above-described regulation is transcriptionally-driven. STRING analysis of downregulated DEPs (FDR <0.1, Log2(fold change) <-0.1) revealed a significantly connected network (expected number of edges: 142; number of edges: 260; PPI enrichment p-value:< 1.0e-16) with overrepresentation of terms related to immunity, myelin, mitochondria and ribosome. The analysis of downregulated DEGs (FDR <0.1, DESeq2) revealed a significantly connected network (expected number of edges:320 ; number of edges: 1482; PPI enrichment p-value:< 1.0e^-16^) with prominent overrepresentation of immunity-related terms and an interesting overrepresentation of the specific WiKI pathway WP3625 Tyrobp causal network in microglia. This network was identified by causal analysis of AD-driving mutations [42]. These data confirm the analysis of GO term overepresentation and indicate that antagonism of miR-29 family dampens the inflammatory reaction induced by the injection via a transcriptional mechanism and reduces myelin, a major plasticity brake [43, 44], via transcript-independent mechanisms.

### Elevated miR-29a levels in the hippocampus of adult mice result in impaired memory retention

To reveal whether miR-29a levels are quantitatively related to memory retention, we asked whether enhancing miR-29a levels in the dorsal hippocampus of adult mice could result in impaired retention. Before conditioning, we injected animals with a synthetic miR-29a mimic (mim29a) and then we tested defensive freezing responses using the same TFC protocol employed in the anti-miR29a experiments (Figure 5A). Mim29a treatment resulted in an upregulation of miR-29a expression and a concurrent downregulation of *Dnmt3a* levels in the dorsal (Figure 5B-C) but not in the ventral hippocampus (Supplementary Figure 8) confirming the spatial selectivity and biological activity of the treatment. At the behavioral level, increasing miR-29a levels affected the trace stability but not the CS response. Indeed, during the extinction protocol (Early Extinction) mim29a mice showed an impaired trace fear retention compared to the control group (Figure 5F). A trend toward a reduced response to the trace interval was also evident during both the training and recall phases (Figure 5D and Supplementary Figure 9A-C). As observed also with anti-miR29a, mim29a treated mice displayed a normal extinction profile (Figure 5F-G). The bidirectional impact of miR-29a on memory strength during the extinction was also evident for contextual memories. In particular, mice receiving miR-29a mimics displayed a strong deficit in the recall of contextual fear memories (Figure 5E) as compared to controls. The involvement of miR-29a in the maintenance of hippocampal fear memory stability was also confirmed by the presence of an inverse correlation between miR29a expression levels and the percentage of freezing behavior exhibited during early extinction and contextual fear recall (Figure 6A-E). Overall, these findings support the hypothesis that heightened miR-29a levels contribute to the compromised stability of hippocampal fear memories [16, 32, 45].

**Figure 5:**
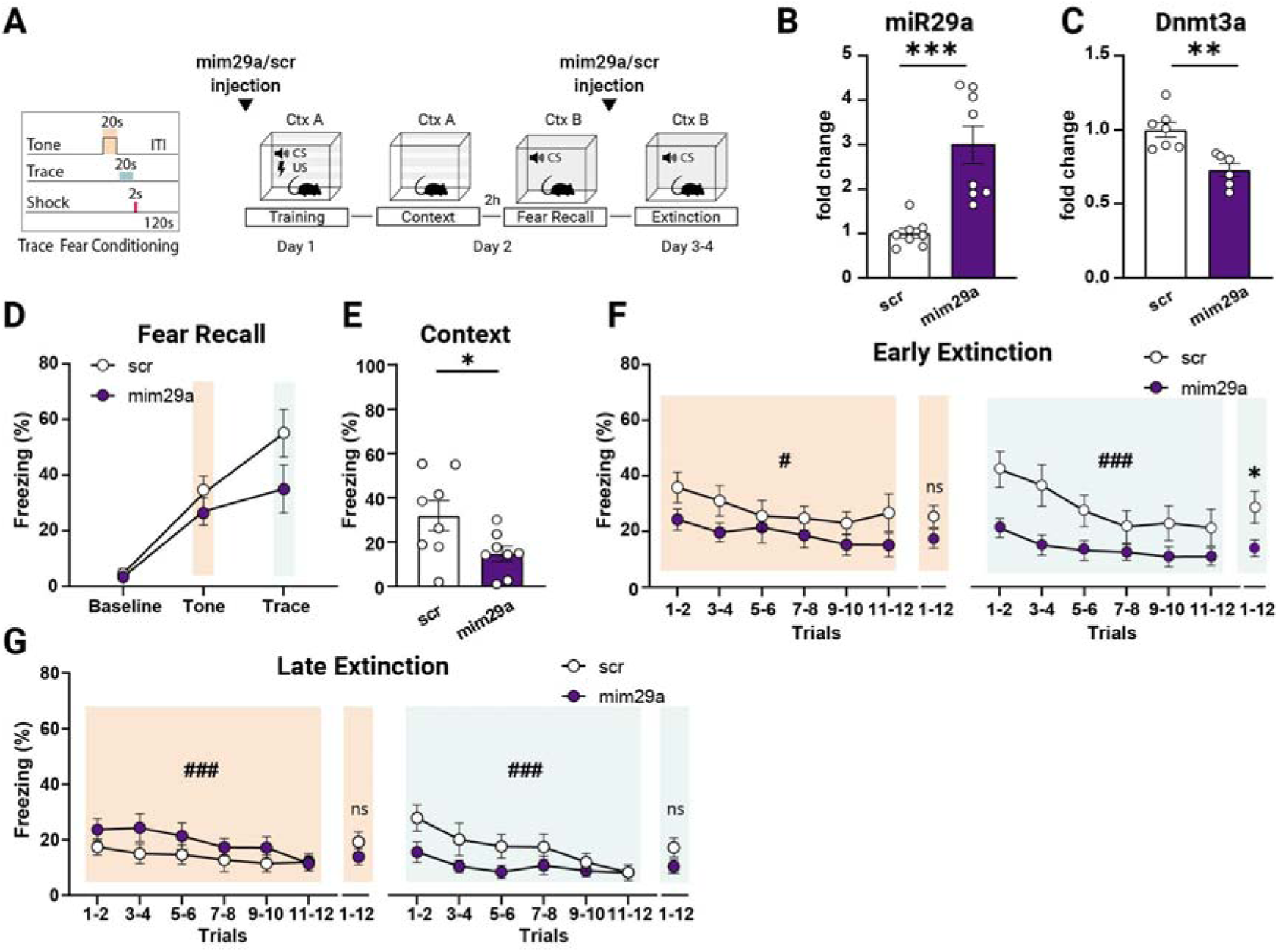
Hippocampal miR-29a upregulation impaired memory retention. **(A)** Diagram showing the TFC paradigm and timeline. To guarantee elevated miR-29a levels during all the sessions of the TFC test, mice were injected 24 hours before the start of the behavioral tests and again 24 hours before the start of the extinction protocol. **(B-C)** Effects of miR-29a mimic treatment on **(B)** miR-29a expression levels (fold change values normalized to scr treated animals; scr: N=8, mim29a N=8; Mann-Whitney U test: U = 0, p *<* 0.001) and **(C)** *Dnmt3a* expression levels (fold change values normalized to scr treated animals; scr: N = 7, mim29a N = 6; Mann-Whitney U test: U = 0, p *<* 0.01). The lower sample size for *Dnmt3a* reflects insufficient RNA in some samples; all available data were included. **(D)** No treatment differences were observed at baseline or during the final two-tone and trace trials of the TFC fear recall session (Two-way ANOVA, test interval × treatment interaction F(1, 42) = 3.52, p = 0.07; main effect of test interval F(2, 42) = 23.89, p *<* 0.0001; main effect of treatment, F(2, 42) = 1.30, p = 0.28). **(E)** MiR-29a levels increase resulted in impaired contextual memory recall (Mann-Whitney U test: U = 12, p *<* 0.05, scr: N = 8, miR-29a mimic N = 8). **(F)** Percentage of freezing during Early extinction. Mim29a treated mice showed lower fear retention compared to the control group during the trace (Two-way RM ANOVA, main effect of treatment F(1, 19) = 4.475, p *<* 0.05; main effect of trials F(3.26, 62.02) = 9.365, p *<* 0.0001; trials × treatment interaction F(5, 95) = 0.77, p = 0.57) but not during the tone interval (Two-way RM ANOVA, main effect of treatment F(1, 14) = 2.30, p = 0.15; main effect of trials F (3.21, 44.97) = 3.80, p *<* 0.05; trials × treatment interaction F(5, 70) = 0.65, p = 0.66). **(G)** During the Late extinction, mim29a treated mice showed no significant differences in the percentage of freezing compared with controls (Tone: Two-way RM ANOVA, main effect of treatment F(1, 14) = 1.33, p = 0.27; main effect of trials F(3.63, 50.83) = 6.41, p *<* 0.001; trials × treatment interaction F(5, 70) = 1.46, p = 0.21; Trace: Two-way RM ANOVA, main effect of treatment F(1,14)) = 2.54, p = 0.13; main effect of trials F(3.35, 46.88) = 9.09, p *<* 0.0001; trials × treatment interaction F(5, 70) = 2.14, p = 0.07). scr: N = 8, mim29a N = 8; *indicates the main effect of treatment, #indicates the main effect of trials.

**Figure 6:**
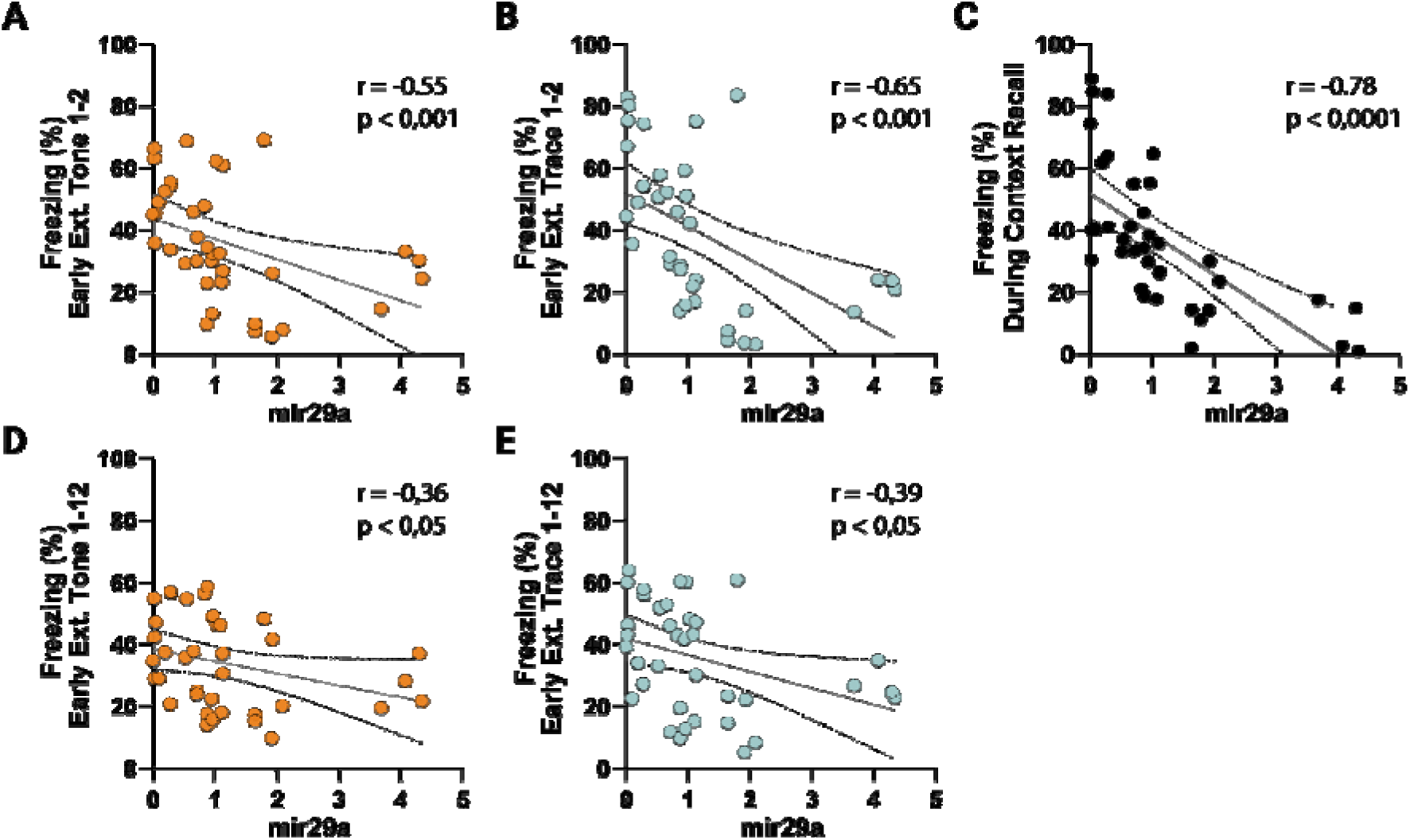
Behavioral correlation with miR-29a levels post LNA and mimic injection. **(A)** During early extinction miR-29a levels negatively correlate with freezing responses at the first two tone and **(B)** trace presentations (Spearman correlation). **(C)** MiR-29a levels negatively correlate with freezing responses during contextual fear recall (Spearman correlation). **(D)** During early extinction miR-29 levels negatively correlate with freezing responses to the tone across all trials **(E)** and with freezing responses during the trace period across all trials (Spearman correlation).

## Discussion

Regulation of the miR-29 family is characterized by a striking degree of evolutionary conservation. Previous studies have demonstrated that miR-29 isoforms exhibit an age-associated upregulation across diverse species and tissues [11, 12, 46]. Furthermore, miR-29a plays a pivotal role in governing age-dependent processes such as neuronal maturation and iron accumulation within the brain [4, 8]. In the visual cortex, miR-29a is the microRNA with the most significant upregulation during the visual critical period [10, 15] [15]. Our data confirmed this age-dependent upregulation of endogenous miR-29a expression in the dorsal hippocampus of wild-type mice, revealing a tenfold increase between P10 and P60. Importantly, a large-scale miRNA association study in human postmortem brain samples identified miR-29a among the miRNAs most strongly associated with cognitive trajectories, with higher miR-29a levels emerging as the second most significant correlate of accelerated cognitive decline (Wingo et al. 2022). Together, these observations suggest that elevated miR-29 levels may be linked to reduced plasticity and cognitive performance. To directly investigate the functional role of miR-29a in memory, we bidirectionally modulated its levels in the dorsal hippocampus of adult mice. Downregulation of miR-29a enhanced hippocampus-dependent memory strength, whereas miR-29a overexpression impaired memory performance, resembling aspects of age-associated cognitive decline. It is important to note that the anti-miR-29a antagomir used in our experiments also targets miR-29c. Given that miR-29a and miR-29c differ by only a single nucleotide and share the same seed sequence, they are expected to regulate largely overlapping target genes; thus, the observed effects may reflect combined modulation of both isoforms. The tight relationship between miR-29 levels and memory is emphasized by the negative correlation between miR-29a levels and the percentage of freezing observed during early extinction and contextual fear recall. Conversely, levels of *Dnmt3a* exhibited a positive correlation with the percentage of freezing exhibited in the initial stages of trace extinction. This finding lends support to the specific regulatory role of miR-29 in memory persistence. Recent work shows that miR-29a downregulation in the hippocampus of 5XFAD mice results in amelioration of memory performances, reduction of beta-amyloid deposition, and lower microglia and astrocytes activation [16]. Collectively, these studies support a model in which miR-29a acts as a key regulator of age-associated molecular pathways that constrain plasticity and memory maintenance.

### miR-29 sculpts the epigenetic landscape of adult hippocampus

Epigenetic mechanisms have been increasingly recognized as important regulators of cognitive functions through the modulation of transcriptional responses [47, 48]. Since we found a strong negative correlation between miR-29a and *Dnmt3a* levels, we decided to explore the profile of methylation of anti-miR29a treated mice. DNA methylation is an essential epigenetic mechanism for various cellular processes and has emerged as a crucial regulator of memory consolidation [49, 50]. anti-miR29a treatment in the adult visual cortex reactivates ocular dominance plasticity in adult mice and upregulates genes associated with the extracellular matrix and epigenetic remodeling, however whether manipulations of miR-29 levels could result in global changes of DNA methylation was utterly unexplored. Contrary to what is observed for miR-29 levels, the degree of DNA methylation decreases with age in many tissues, including the nervous system [51, 52]. In particular, aging has been associated with a reduction in the expression of the DNA methyltransferase *Dnmt3a2* in the hippocampus, and restoring *Dnmt3a2* levels has been shown to rescue cognitive functions affected by aging [32]. It has also been reported that hippocampal *Dnmt3a2* levels determine cognitive abilities in both young adult and aged mice [53–56]. Kupke et al, however, showed that Dnmt3a1 regulates hippocampus-dependent memory via Nrp1 [57].

Furthermore, DNA methylation induced selectively within neuronal ensembles has been proposed as a mechanism for stabilizing engrams during consolidation, supporting successful memory retrieval [58]. Consistent with this framework, inhibition of miR-29a led to a robust increase in CpG methylation across the hippocampal genome, in parallel with increased *Dnmt3a* expression. Importantly, these methylation changes were substantially blocked by *Dnmt3a* downregulation, demonstrating that a major component of the epigenetic remodeling induced by miR-29a antagonism depends on DNMT3a activity. A direct comparison of DMCpGs between anti-miR29a alone and the combined anti-miR29a + anti-DNMT3a treatment revealed a highly non-random distribution of concordant and discordant CpGs, supporting a structured and pathway-specific epigenetic response. Concordant hypermethylated CpGs, those remaining hypermethylated even after DNMT3a inhibition, were enriched for synaptic processes, including vesicle-mediated transport at the synapse. In contrast, discordant CpGs, hypermethylated after anti-miR29a but hypomethylated when DNMT3a was simultaneously inhibited, were enriched for processes such as regulation of membrane potential, nervous system development, forebrain development, and cell junction assembly. These findings indicate that miR-29a inhibition affects multiple neuronal pathways, with a subset of methylation changes being DNMT3a-dependent and others likely mediated through additional mechanisms. Global CH methylation was also significantly altered across treatment groups, further supporting a broad DNMT3a contribution to the epigenomic remodeling triggered by miR-29a antagonism. Gene Ontology and KEGG pathway analyses reinforced the functional relevance of these methylation changes. Hypermethylated CpGs following anti-miR29a treatment were enriched in pathways related to synaptic signaling, extracellular matrix organization, kinase activity, and notably the glutamatergic synapse pathway. Mapping methylation changes onto the KEGG glutamatergic synapse pathway revealed widespread hypermethylation across synaptic genes, suggesting that miR-29a inhibition engages epigenetic programs closely linked to hippocampal-dependent plasticity and memory persistence. While promoter methylation is often associated with transcriptional repression, gene body methylation is positively correlated with gene expression [59, 60], indicating that the observed hypermethylation may also facilitate transcriptional activation of plasticity-related genes rather than suppress them. To explore the functional consequences of this epigenetic remodeling, we performed RNA sequencing and mass-spectrometry–based proteomics. Both datasets converged on a coherent molecular signature. Upregulated transcripts and proteins were strongly enriched for synaptic and plasticity-related pathways, with glutamatergic transmission (KEGG mmu04724) standing out prominently. Glutamate receptors and transporters, including NMDA receptor subunits, were upregulated at both transcript and protein levels. Network analysis revealed highly interconnected clusters composed predominantly of synaptic proteins and RNA-binding proteins, indicating a coordinated and transcriptionally driven enhancement of synaptic machinery. The enrichment of miR-29 binding sites among upregulated genes confirmed effective antagonism of the miR-29 family and supports the view that these transcriptional changes are directly linked to miR-29 inhibition.

Conversely, downregulated transcripts were strongly enriched for genes expressed in microglia and other immune-related cell types, including components of the Tyrobp causal network implicated in Alzheimer’s disease. Lysosomal pathways were also significantly reduced. At the protein level, downregulated groups were enriched for oligodendrocyte-specific and myelin-associated proteins. Together, these findings indicate that miR-29 antagonism dampens inflammatory and microglial activation through transcriptional mechanisms while reducing myelin-related proteins through mechanisms that may be partially transcription-independent. Given that myelin constitutes a well-established brake on plasticity [43, 44], its reduction may further contribute to the enhanced plastic potential observed after miR-29 inhibition. Overall, these results delineate a coordinated molecular program in which miR-29a antagonism promotes DNMT3a-dependent epigenetic remodeling, enhances glutamatergic synaptic function, suppresses inflammatory signaling, and reduces structural constraints on plasticity. This integrated regulation could be further complemented by studies aimed at identifying the specific cell types involved in miR-29a action, thereby providing a more refined mechanistic framework linking miR-29a levels to memory persistence. Future studies examining memory performance across developmental stages characterized by distinct endogenous miR-29a levels will further clarify how this regulatory axis contributes to age-dependent changes in cognitive function.

## Resource availability

### Lead contact

Further information and requests for resources and reagents should be directed to the lead contact, Tommaso Pizzorusso.

### Materials availability

This study did not generate new unique reagents.

### Data and code availability

● All data reported in the main text or supplemental information are available from the lead contact upon request.
● All original code is also available from the lead contact upon request.
● Any additional information required to reanalyze the data reported in this paper is available from the lead contact upon request.

## Supporting information

All supplemental figures

## Acknowledgments

We gratefully acknowledge NVIDIA Corporation’s support with the Jetson AGX Xavier Developer Kit donation for this research. European Union-Next Generation EU, Mission 4 Component 1 CUP E53C24001460006, project TNE-NEUROBRIDGE and PRIN2022 20228RMXBE to T.P.

## Author contributions

Conceptualization: AV, AC, TP

Methodology: AV, EP, CG, RM, SB, PT, AC, TP

Formal analysis: AV, CG, SB, PT, AC,

Investigation: AV,CG,EP, SB, PT,

Resources: AC, TP, RM

Data curation: AV, CG, SB, AC

Writing original draft: AV, AC, TP

Visualization: AV, CG, SB, AC

Supervision: TP, AC

Project administration: TP, AC

Funding acquisition: TP, AC

## Experimental Procedures

### Animals

All experiments utilized the C57BL/6J strain. Animals were maintained at 22°C under a 12-h light–dark cycle (average illumination levels of 1.2 cd/m2) and housed in standard cages according to current regulations about animal welfare. Food (4RF25 GLP Certificate, Mucedola) and water were available ad libitum. All the experiments were carried out according to the directives of the European Community Council (2011/63/EU) and approved by the Italian Ministry of Health (621/2020-PR). All tissue explants were performed at the same time of the day (10–12 AM, Central European Time (CET, Italy)). Both male and female mice were included in the study, and no significant sex × treatment interactions were detected (Supplementary Table 1).

### Stereotaxic intracranial injections

Mice were anesthetized with isoflurane (3% induction and 1.5% maintenance) and secured in a stereotaxic frame (Sutter Instrument, Novato, CA, USA) using ear bars. Prilocaine was used as a local anesthetic for the acoustic meatus. Body temperature was maintained at 37° C using a heating pad. The eyes were treated with a dexamethasone-based ophthalmic ointment (Tobradex, Alcon Novartis) to prevent cataract formation and keep the cornea moist. Respiration rate and pedal reflex were checked periodically to maintain an optimal level of anesthesia. Prior to scalp removal, a subcutaneous local injection of lidocaine (2%) was performed. After exposing the skull, small burr holes were made using a surgical drill. In separate experiments, mice received either LNA oligonucleotide complementary to miR-29a (miR-29a-3p miRCURY LNA, QIAGEN, Cat. No. 339121, 50 µM) or a synthetic miR-29a mimic (mirVana™ miRNA miR-29a-3p Mimic, Thermo Fisher Scientific, Cat. No. 4464066, 100 µM). The corresponding scrambled oligonucleotides were used as controls (Negative Control miRCURY LNA miRNA Mimic, QIAGEN, Cat. No. YM00479902, 50 µM; mirVana™ miRNA Mimic Negative Control #1, Thermo Fisher Scientific, Cat. No. 4464058, 100 µM). To assess the contribution of *Dnmt3a* to the effects mediated by miR-29a, a GapmeR antisense oligonucleotide targeting murine *Dnmt3a* was injected. In these experiments, the GapmeR anti-*Dnmt3a* in combination with the miR-29a LNA was delivered into one hippocampus, while the contralateral hippocampus received the corresponding control oligonucleotides. Additional control conditions included injections of GapmeR anti-*Dnmt3a* alone or the respective control oligonucleotides. Injection in the dorsal hippocampus was done using the following stereotaxic coordinates: –2 mm anteroposterior, ±1.5 mm mediolateral, –1.5, –1.7 mm dorsoventral from bregma. A total of 2 µl of solution per hemisphere was injected, 1µl for each coordinate using a Hamilton syringe. The injection needle was left in place for an additional 60 s to allow the fluid to diffuse. The skin was subsequently sutured, and a physiological solution was injected subcutaneously to prevent dehydration. At the end of the surgical procedure, mice were left in a heated cage for recovery. Once fully awake, they were returned to their home cage. Paracetamol was administered in water ad libitum for two days. Due to the inherent stability of LNA oligonucleotides, a single injection was administered to mice before the beginning of the behavioral test. In contrast, the miR-29a mimic exhibited faster degradation within a specific three-day timeframe. To maintain the levels of miR-29a consistently elevated during all the sessions of the behavioral test, mice received injections every two days, following the protocol reported in Napoli et al., 2020.

### Synthesis and selection of GapmeR Antisense Oligonucleotides

Three LNA GapmeR antisense oligonucleotides targeting the sequence (AGGGTACTGGCCGCCTCTTCTTTGAGTTCTACCGCCTCCTGCATGATGCGCGGCCCAAGGAGGGAGATGATCGCCCCTTCTTCTGGCTCTTTGAGAATGTGGTGGCCATGGGCGTTAGTGACAAGAGGGACATCTCGCGATTTCTTGAGGTATAGACCGAGACCTTGGTTTGGCCAGCTCACTAATGGCTTCTACCTGGGACTGCTGCTTCGTCCCTGTCTTGTCTGCATTGCGGAGCTGGGGGATTGGAGCTGGGGACTGGTGGCTTCTCTTTGCAAGGGACAGCTTGAGGAAGATTTTCCATGTAGAGAGGAAGCAGTGCTAAAGACCCACCTAGGAAGAAAGTTCTCTTGTTCAAAGAGGTGTGGTGGTCATCATCAAACAGATGGACTGGGGCCAGCCCAGTTTTTCTGTGGAGAACTCCAAAATCAGTTTTTAAAATCATTCTCTGACTAGAATGGCCTGTGCTCACTCTCTGGCACCCTTGTGGGTGTTTTGTGATACCTGAGAGAAAATCATGGCTTATCTTTTTGCTTTCATTTCTTGTTATGTGAGCCCCAGTTTAGCACTGAGCCATGTACAAGCTCAATCATAGGGGATTGCTGCTGCCCAAGGCATTCTTTTCTTTTCTTTTCTTTTCTTTTCTTTTCTTTTCTTTTCTTTTCTTTCTCTCTCTCTCTCTTTCTTTCTTTCTTCCTTTCTTTCTTTCTTTTTGATTTATTTATTATTATATATAAGTAGCTGACTTCAGACACACCAGAAGAGGGTGTCAGATCTCATGGGTGGTTGTGAGCCACCATGTGGTTGCTGGGATTTGAACTCAGGACCCTCGGAAGAGCAGTCAATGCCCTTAACTGCTGAGCCATCTCTCCAGCCCTGCCCAAGGCATTCTTGTGGTAGGCTGTCAGCTTATAGTCCTGTCAGCCTACGCTCAATAATAACCTCAGATTGTAAATGTGAGGACTTAGTACACAAACAGGCTTGCTCCGGTAAGGTCTTATGTGGTCATCAGCCTTGTGGCTATTACAAGAATGAACCTCACTTTAAGCACACAAGAGTCATTCATGGTACTACTGGAATAATAACAAAGTTCAGTTGGATTGTGCTGGCTGTCCAGAATGTTCTAGCCAGCCGGGCGTGGTGGCTCACGCCTTTAATCCCAGCACTTGGGAGGCAGAGGCAGGTGGATTTCTGAGTTCGAAGCCACCCGGTCTACAAAGTGAGTTCCAGGACAGCCAGGGCTATACAGAGAAACCCTGTCTTGAAAAATAAAAATTAAAAAAAAAAAAAAAGAATGTTCTAGCCAGCACATGTGGGATGGATTTAAGTTTGGGTCTTCAGGTTATTCCGGTGTTTGGCTTCCTACGGAGGAAGTTCCTCTGGGGAGTTAAGACTCTTGGCCAGCCCAGTTGCCCAGCAGTGTTTAATTAATACTTCCTCCTTGGTCATCTTGAAACCATCTCCTATTTTACAG) of the murine *Dnmt3a* transcript were designed and synthesized by QIAGEN (Germany). A non-targeting control GapmeR was also designed and synthesized by the manufacturer. All GapmeRs contained LNA modifications and a phosphorothioate backbone to enhance nuclease resistance and promote RNase H–mediated degradation of the target RNA, according to the manufacturer’s specifications. To determine the most effective GapmeR in our conditions, adult C57BL/6J mice were anesthetized with isoflurane and placed in a stereotaxic frame. GapmeRs were diluted in sterile PBS and injected into the visual cortex at a final dose of 100 pmol in a total volume of 1 µL per injection site using a Hamilton syringe. The control GapmeR was injected into the contralateral visual cortex of the same animal. Seven days after injection, mice were euthanized and the cortices were rapidly dissected on ice. Tissue samples were snap-frozen in liquid nitrogen and stored at −80 °C until further processing. RNA extraction, reverse transcription and Quantitative real-time PCR for murine *Dnmt3a* were performed as described below (see section: RNA extraction). GapmeR efficacy was determined by comparing *Dnmt3a* mRNA levels in treated cortices relative to those injected with the control GapmeR. Among the three candidates tested, the GapmeR producing the greatest reduction in *Dnmt3a* transcript levels was selected for subsequent experiments (anti-*Dnmt3a*).

### Trace fear conditioning and extinction protocol

Mice were subjected to a trace fear conditioning and extinction procedure using a custom-made PVC/acrylic apparatus consisting of three radially arranged arms converging at a central junction. Each arm measured 50 cm in length, 15 cm in width, and 21Lcm in height, and was covered by a clear plastic lid to prevent escape and allow behavioral observation. The floors consisted of electrifiable grids of parallel stainless steel bars, permitting delivery of aversive foot shock stimuli. A circular speaker of 5 cm diameter was mounted centrally in the lid of each arm to deliver precisely controlled auditory cues (conditioned stimuli). The arms differed in wall and floor textures to provide distinct contextual cues, allowing for contextual discrimination between conditioning and extinction environments.. During the test days, the mice were transported in their home cages to a room adjacent to the testing room and left for 2 h before behavioral testing. We used two different contexts: In context A, the walls had white plastic circles and the floor was completely black; in context B, the walls had white vertical plastic strips and a white floor. Both chambers were covered with transparent plexiglass lids with a loudspeaker in its center point. The shock grid on the floor was made of stainless steel. Only the grid of context A was electrified by a shock generator (World Precision Instruments, Sarasota, FL) and guided by an Arduino Uno for CS and US parameter control and footshock delivery. Mice behavior was recorded by a camera controlled by the EthoVision XT 8 software (Noldus Information Technology, The Netherlands). The apparatus was cleaned before and after each animal with 70% ethanol or 1% acetic acid for context A and context B respectively, since the mice may associate the smell with the context. After each session, mice were housed separately until the end of the test to avoid possible observational fear learning. During the Training phase of the test (Day 1), mice were allowed to explore the conditioning chamber (Context A) for 2 min (baseline) before receiving five conditioning trials. Each trial consisted of a 20 s pure tone (80 dB, 2900 Hz) and a 2 s shock (0.6 mA) separated by a 20 s stimulus-free trace interval. The intertrial interval (ITI) was 120 s. Mice were removed from the chamber 120 s after the last trial. Twenty-four hours later (Day 2), mice were placed back in the conditioning chamber (Context A) for a 5 min context test. The same day (Day2) after 2 h from the context test, mice were placed in a different context (Context B) for a Recall test consisting of a 2 min baseline period followed by four 20 s tone presentations separated by a 20 s stimulus-free trace interval with 120 s ITI. On day 3 (Early Extinction) and day 4 (Late Extinction), conditioned mice were subjected to the extinction training in context B during which they received twelve 20 s tone presentations separated by 20 s stimulus-free trace interval presentations with 120 s ITI each day.

### Analysis of Freezing Behavior

Recorded videos were manually scored for freezing behavior by two separate experimenters blind to treatment conditions. Mice were considered to be freezing if no movement was detected for 2 s (defined as the complete absence of movement except for respiratory movements). Evoked freezing behavior was analyzed by calculating the percentage of time an animal spent freezing during a given phase of the test. During fear extinction, averages were calculated by pooling freezing across 2 CS presentations if not indicated otherwise.

### RNA extraction and quantification (qPCR)

In all experiments, mice were euthanized and brains were rapidly removed and placed on ice. The dorsal and ventral hippocampus were dissected as reported in Jaszczyk et al. 2022. Briefly, the two hemispheres were separated and the brain was rotated to expose the medial surface. The hippocampus was carefully exposed by gentle mechanical dissection, detached from the surrounding tissue, and cleaned of residual white matter. The isolated hippocampus was then divided into three portions—dorsal, intermediate, and ventral—based on anatomical landmarks and millimeter measurements. Tissue samples were snap-frozen in liquid nitrogen and stored at −80 °C until further processing. Tissue samples were homogenized in the cell disruption buffer (Ambion). RNA was extracted by the addiction of Phenol/guanidine-based QIAzol Lysis Reagent (Qiagen, cat. no. 79306). Chloroform was added, and the samples were shaken for 15 s. The samples were left at 20–24°C for 3 min and then centrifuged (12,000 g, 20 min, 4°C). The upper phase aqueous solution, containing RNA, was collected in a fresh tube, and the RNA was precipitated by the addition of isopropanol. Samples were mixed by vortexing, left at 20–24°C for 15 min, and then centrifuged (12,000 g, 20 min, 4°C). The supernatant was discarded, and the RNA pellet was washed in 75% ethanol by centrifugation (7500 g, 10 min, 4°C). The supernatant was discarded, and the pellet was left to dry for at least 15 min; then, it was resuspended in RNAse-free water. Total RNA concentrations were determined by NanoDrop Spectrophotometer (Thermo Scientific 2000 C). RNA quality was analyzed through a gel running (1% agarose). Total RNA was reverse transcribed using the QuantiTeck Reverse Transcription Kit (Qiagen, cat. no. 205311), and miRNAs were reverse transcribed using the TaqMan MicroRNA reverse transcription kit (Thermo Fisher, cat. no. 4366596). Gene expression was analyzed by real-time PCR (Step one, Applied Biosystems). TaqMan inventoried assays were used for miR-29a-3p (assay ID: 002112), sno234 (assay ID:001234), and *Dnmt3a* (assay ID: Mm00432881_m1). TaqMan assay was used for glyceraldehyde 3-phosphate dehydrogenase (Gapdh), GAPDH probe ATCCCAGAGCTGAACGG, GADPH forward CAAGGCTGTGGGCAAGGT, and GADPH reverse GGCCATGCCAGTGAGCTT. Quantitative values for cDNA amplification were calculated from the threshold cycle number (Ct) obtained during the exponential growth of the PCR products. The threshold was set automatically by the Step One software. Expression levels were normalized using *Gapdh or sno234* as housekeeping. Relative gene expression was calculated using the ΔΔCt method.

### DNA extraction and validation

For the methylation analysis, we administered LNA-anti-miR29a to one hemisphere of seven mice, while the other hemisphere received an injection of the scrambled sequence, serving as an internal control. To mitigate potential lateralization effects, we randomized the side of LNA injection. Seven days after the injection, tissues were collected and samples were stored in an ultralow freezer (−80°C) until DNA extraction. DNA was isolated from snap-frozen hippocampi using the QIAamp DNA Micro Kit (Qiagen) following the manufacturer’s protocol. DNA concentration was measured with NanoDrop 2000 (Thermo Scientific), using 1 µl of input DNA. Then, dsDNA concentration was quantified using Quant-iT Picogreen dsDNA Assay (Thermo Scientific), and the distribution of DNA fragments was assessed by TapeStation 4200 (Agilent).

### Reduced Representation Bisulfite Sequencing (RRBS)

RRBS libraries were produced by Ovation RRBS Methyl-Seq with the TrueMethyl oxBS kit (Tecan, Redwood City, CA, USA) according to the manufacturer’s instructions. Briefly, 100 ng of genomic DNA was digested for 1h at 37°C with the methylation-insensitive restriction enzyme MpsI. Then, fragments were ligated to methylated adapters and treated with bisulfite to convert unmethylated cytosine into uracil. PCR amplification was then performed to obtain the final DNA library. The resulting libraries were sequenced on the Illumina NovaSeq6000 with 100 bp single-end sequencing with an average of 60 million reads per sample.

### RRBS data analysis

The sequencing reads were trimmed to remove the adapter and low-quality bases with Trim Galore! (www.bioinformatics.babraham.ac.uk). Then trimmed reads were aligned to the mouse reference genome mm39 with BSMAPz (github.com/zyndagj). Using the Python script methratio.py in BSMAPz, methylation ratios were extracted from the mapping output. Methylation analysis was performed using the R package methylKit (version 1.20.0) (Akalin et al., 2012). Only CpGs covered by at least 10 reads and present in every sample were retained for downstream analysis. The methylation ratio of each site was calculated by dividing the number of reads called “C” by the total number of reads called either “C” or “T” at the specific site. The methylation score for each CpG site is represented as a β-value which ranges between 0 (unmethylated) and 1 (fully methylated). Differentially methylated CpGs (DMCs) were detected using a logistic regression model based on a Chi-square test. A false discovery rate (FDR) q value threshold of < 0.05 and methylation difference of ±5% between treated and control groups was used to identify significant DMCs. The promoter was defined as the region ±1kb from the transcription start site. CpG sites located on the X and Y chromosomes were excluded from the methylation analysis due to a strong sex-associated effect, which represented the main source of variability in the dataset. All downstream analyses were therefore restricted to autosomal CpGs. Differential methylation analysis was performed separately for each treatment group (anti-miR29a, anti-DNMT3a and combined anti-miR29a + anti-DNMT3a) compared to control samples using methylKit. Multivariate analysis was performed using partial least squares discriminant analysis (PLS–DA) in mixOmics R package to assess sample clustering based on methylation profiles; the first two components were used for visualization. To evaluate concordance of methylation changes between treatment conditions, DMCpGs identified in the combined treatment were compared to those identified in the anti-miR29a treatment alone. CpGs were classified into four quadrants based on the direction of methylation change. A χ² test was performed to assess deviation from a uniform distribution of concordant versus discordant CpGs. Non-CpG (CH) methylation levels were also quantified. Per-animal Δ methylation values (treated minus control hemisphere) were calculated and compared across treatment groups using Welch’s ANOVA followed by Bonferroni-corrected pairwise t-tests. Pathway analysis was performed using the web tools WebGestalt (www.webgestalt.org). Pathway visualization was performed using the R package Pathview to map methylation changes onto KEGG pathways. The KEGG pathway ID mmu04724 (Glutamatergic synapse) was specifically examined to visualize gene-level methylation changes following anti-miR29a treatment.

### Sample preparation for proteomics analysis

Before lysis, a lysis buffer was added to each sample to reach final concentrations of 4% SDS, 100 mM HEPES (pH 8.5), and 50 mM DTT. Samples were then boiled at 95°C for 7 min and sonicated using a tweeter. Reduction was followed by alkylation with 200 mM iodoacetamide (IAA, final concentration 15 mM) for 30 min at room temperature in the dark. Samples were acidified with phosphoric acid (final concentration 2.5%), and seven times the sample volume of S-trap binding buffer was added (100 mM TEAB, 90% methanol). Samples were bound on a 96-well S-trap micro plate (Protifi) and washed three times with a binding buffer. Trypsin in 50 mM TEAB pH 8.5 was added to the samples (1 µg per sample) and incubated for 1 h at 47°C. The samples were eluted in three steps with 50 mM TEAB pH 8.5, elution buffer 1 (0.2% formic acid in water) and elution buffer 2 (50% acetonitrile and 0.2% formic acid). The eluates were dried using a speed vacuum centrifuge (Eppendorf Concentrator Plus, Eppendorf AG, Germany) and stored at -20° C. Before analysis, samples were reconstituted in MS Buffer (5% acetonitrile, 95% Milli-Q water, with 0.1% formic acid) and spiked with iRT peptides (Biognosys, Switzerland).

### LC-MS Data independent analysis (DIA)

Peptides were separated in trap/elute mode using the nanoAcquity MClass Ultra-High Performance Liquid Chromatography system (Waters, Waters Corporation, Milford, MA,USA) equipped with a trapping (Waters nanoEase M/Z Symmetry C18, 5μm, 180 μm x 20 mm) and an analytical column (Waters nanoEase M/Z Peptide C18, 1.7μm, 75μm x 250mm). Solvent A was water and 0.1% formic acid, and solvent B was acetonitrile and 0.1% formic acid. 1 µl of the sample (∼1 μg on column) was loaded with a constant flow of solvent A at 5 μl/min onto the trapping column. Trapping time was 6 min. Peptides were eluted via the analytical column with a constant flow of 0.3 μl/min. During the elution, the percentage of solvent B increased in a nonlinear fashion from 0–40% in 120 min. Total run time was 145 min. including equilibration and conditioning. The LC was coupled to an Orbitrap Exploris 480 (Thermo Fisher Scientific, Bremen, Germany) using the Proxeon nanospray source. The peptides were introduced into the mass spectrometer via a Pico-Tip Emitter 360-μm outer diameter × 20-μm inner diameter, 10-μm tip (New Objective) heated at 300 °C, and a spray voltage of 2.2 kV was applied. The capillary temperature was set at 300°C. The radio frequency ion funnel was set to 30%. For DIA data acquisition, full scan mass spectrometry (MS) spectra with mass range 350–1650 m/z were acquired in profile mode in the Orbitrap with resolution of 120,000 FWHM. The default charge state was set to 3+. The filling time was set at a maximum of 60 ms with a limitation of 3 × 10 6 ions. DIA scans were acquired with 40 mass window segments of differing widths across the MS1 mass range. Higher collisional dissociation fragmentation (stepped normalized collision energy; 25, 27.5, and 30%) was applied and MS/MS spectra were acquired with a resolution of 30,000 FWHM with a fixed first mass of 200 m/z after accumulation of 3 × 10 6 ions or after filling time of 35 ms (whichever occurred first). Data were acquired in profile mode. For data acquisition and processing of the raw data Xcalibur 4.3 (Thermo) and Tune version 2.0 were used.

### Proteomic data processing

DIA raw data were analyzed using the directDIA pipeline in Spectronaut v.18 (Biognosys, Switzerland) with BGS settings besides the following parameters: Protein LFQ method = QUANT 2.0, Proteotypicity Filter = Only protein group specific, Major Group Quantity = Median peptide quantity, Minor Group Quantity = Median precursor quantity, Data Filtering = Qvalue, Normalizing strategy = Local Normalization. The data were searched against a UniProt (Mus musculus, v. 160106, 16.748 entries) and a contaminants (247 entries) database. The identifications were filtered to satisfy FDR of 1 % on peptide and protein level. Relative protein quantification was performed in Spectronaut using a pairwise t-test performed at the precursor level followed by multiple testing correction according to Benjamini-Hochberg.

### RNA-Seq library preparation

Sequencing of RNA samples was done using Illumina’s next-generation sequencing methodology . In detail, quality check and quantification of total RNA was done using the Agilent Bioanalyzer 2100 in combination with the RNA 6000 pico kit (Agilent Technologies, 5067-1513). Total RNA library preparation was done by introducing 500 ng total RNA into Illumina’s NEBNext Ultra II directional mRNA (UMI) kit (NEB, E7760S), following the manufacturer’s instructions. The quality and quantity of all libraries were checked using Agilent’s Bioanalyzer 2100 and DNA 7500 kit (Agilent Technologies, 5067-1506).RNAseq analysis. All libraries were sequenced on a NovaSeq6000 SP 100 cycles v1.5. Corresponding UMI file was used to add UMI sequences to the reads names of each sample R1 file using UMI-tools extract v1.1.1 [73] to enable duplicate removal in the later steps. Trimmomatic v0.36 [74] was applied to the reads to remove adapters and low quality sequences using custom adapter fasta file and the following options adapters_3tools.fa:2:30:10:6 SLIDINGWINDOW:3:25 MINLEN:25. Trimmed reads were then mapped onto the *Mus musculus* GRCm39 genome primary assembly from Ensembl release 111 [75] using STAR v2.7.10a [76] and -outSAMtype BAM SortedByCoordinate - outSAMmultNmax 1 --outSJfilterReads Unique --outSAMstrandField intronMotif --alignIntronMax 100000 --outFilterMismatchNoverLmax 0.04 options to generate sorted bam files. The resulting bam file was indexed using samtools v1.19.2-23-g1a63877 [77] and duplicates were removed from it using UMI-tools dedup v1.1.1 [78]. In order to get gene-level counts for each sample featureCounts v2.0.3 [79] was used with mouse GRCm39 GTF annotation from Ensembl release 111 [80] and -s 2 flag. Counts from individual samples were then combined into a single counts matrix for use in downstream analysis tools.

### Differential expression Analysis

Differential expression analysis was performed using DESeq2 [81]. Exploratory data analysis was performed with PCA to identify possible outliers. Counts were normalized by variance stabilizing transformation (vst) and adaptive fold-change shrinkage (apeglm). Differential expression analysis was performed with default parameters. Since samples are paired, the following design was implemented: design = ∼ Subject + Cond where Cond has two categorical values = “Scramble” and “anti-miR29”.

### Statistical analysis

All the statistical analyses were performed using GraphPad Prism 8. The Shapiro–Wilk test was used to assess the normality of the data. For datasets that deviated significantly from normality, appropriate non-parametric tests were applied. For the two-way mixed ANOVA, this parametric approach was retained because it efficiently assesses main effects and interactions for both within- and between-subject factors. Although the sample sizes were slightly unequal between groups, ANOVA is generally robust to such minor imbalances, particularly when variances are similar. Moreover, the Geisser–Greenhouse correction was applied to account for violations of sphericity, ensuring valid F-tests. Analyses were followed by appropriate post hoc tests. Significance was set at P*<*0.05 for all tests. Error bars represent s.e.m. in all figures.

