## Supplementary material for "MicroRNA-29 acutely regulates Memory Stability, Expression of Synaptic Genes, and DNA Methylation in the Mouse Adult Hippocampus": All supplemental figures

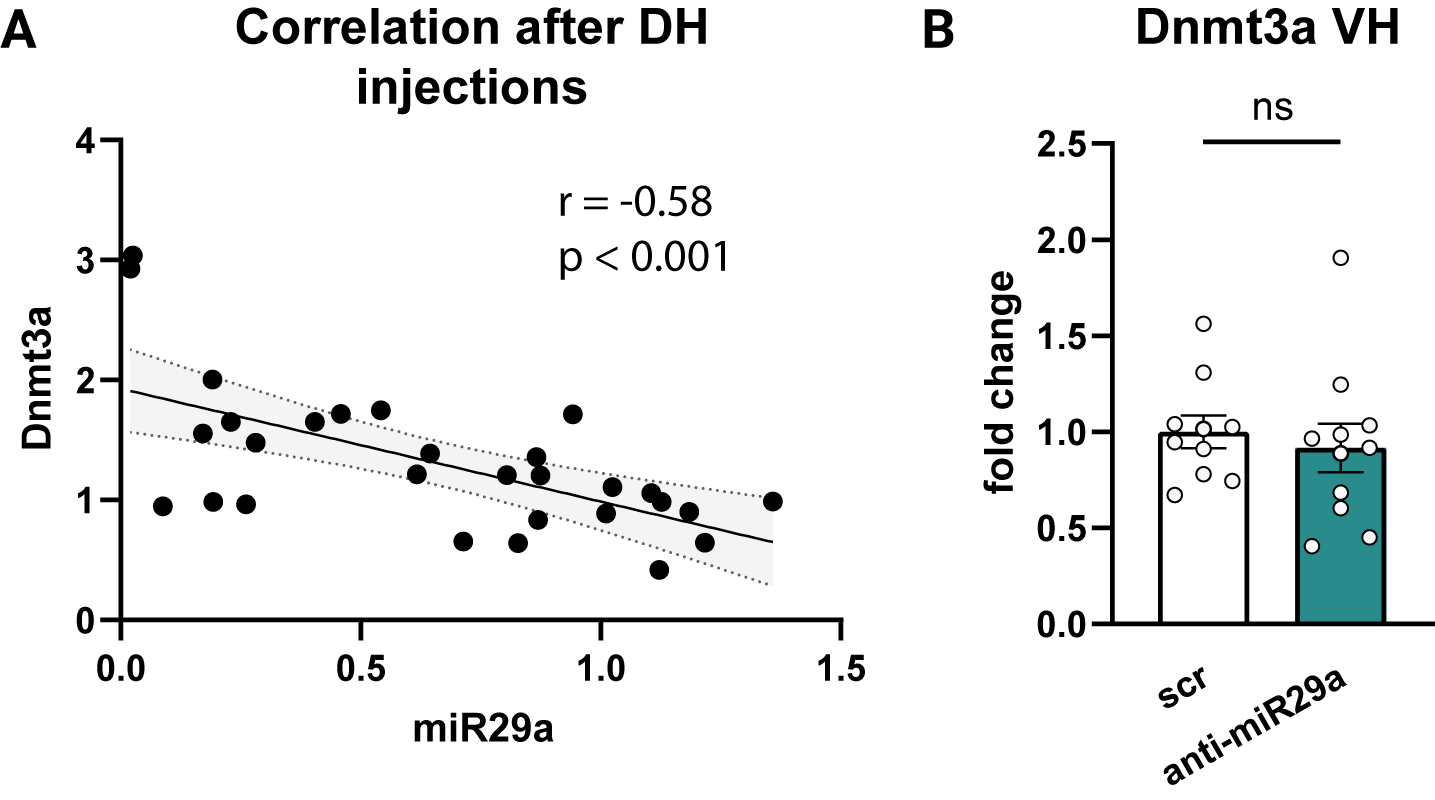

**Supplementary Figure 1. Specificity of anti-miR-29a injection**. **(A)** Spearman’s correlation between *Dnmt3a* and miR-29a levels after the LNA injection. scr: N = 16, anti-miR29a N = 13. **(B)** LNA injection in the dorsal hippocampus did not affect the expression of *Dnmt3a* in the ventral hippocampus confirming the specificity of the injection site within the dorsal hippocampus (Mann-Whitney U test: U = 43, p = 0.4359). scr: N = 10, anti-miR29a N = 11. DH = Dorsal Hippocampus; VH = Ventral Hippocampus.

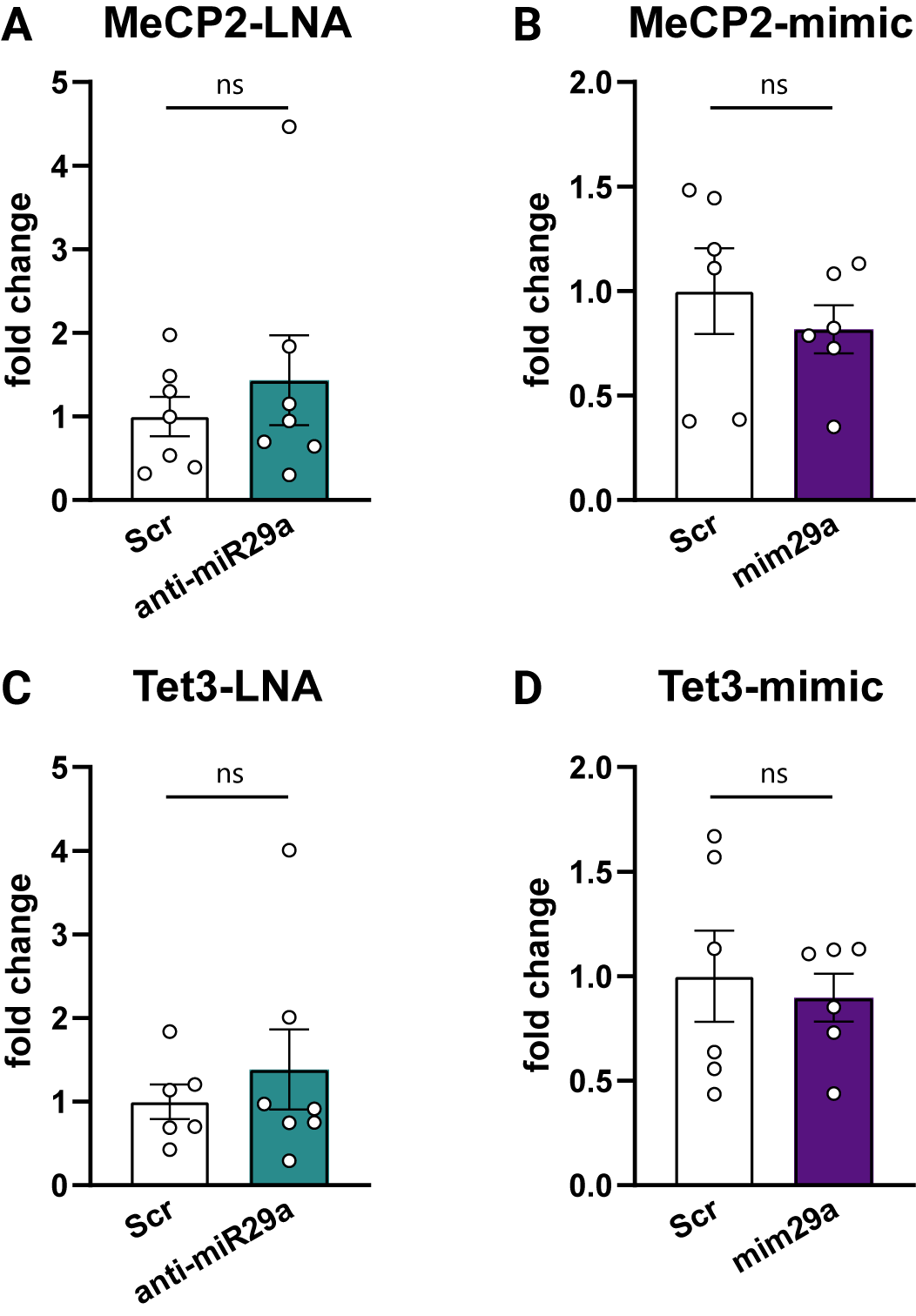

**Supplementary Figure 2. Effects of miR-29a modulation on MeCP2 and Tet3 expressions in the dorsal hippocampus. (A)** MeCP2 expression levels following anti-miR-29a LNA injection compared to scr control (Mann–Whitney U test: U = 23, p = 0.9015; scr N = 7, anti-miR-29a N = 7). One sample from the anti-miR-29a group was excluded as a significant outlier. **(B)** MeCP2 expression following miR-29a mimic administration compared to scr control (Mann–Whitney U test: U = 11, p = 0.3095; scr n = 6, miR-29a mimic n = 6). **(C)** Tet3 expression after anti-miR-29a LNA treatment compared to scr control (Mann–Whitney U test: U = 18, p = 0.7308; scr n = 7, anti-miR-29a n = 7). One sample from the anti-miR-29a group was excluded as a significant outlier. **(D)** Tet3 expression following miR-29a mimic administration compared to scr control (Mann–Whitney U test: U = 16, p = 0.8182; scr n = 6, miR-29a mimic n = 6). All analyses were conducted on samples remaining after the primary experimental analyses.

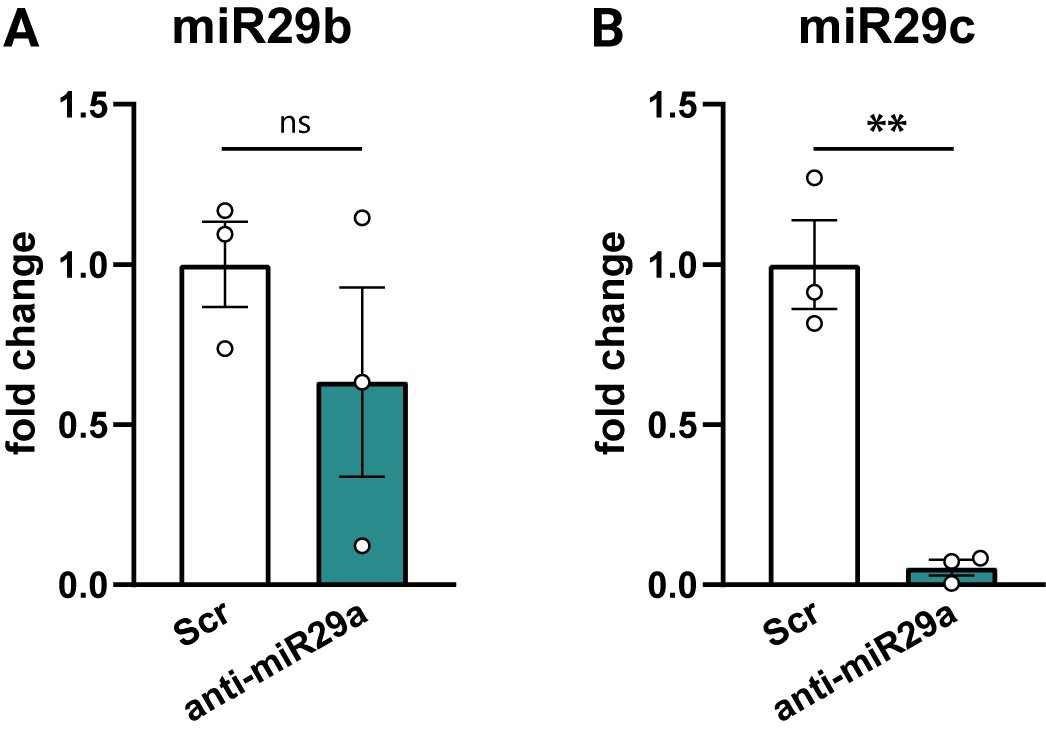

**Supplementary Figure 3. Effects of anti-miR-29a treatment on miR-29b and miR-29c expression. (A)** miR-29b expression was not significantly affected by anti-miR-29a treatment (scr: N = 3; anti-miR-29a: N = 3; unpaired t-test: p = 0.321; Mann–Whitney U test: U = 2, p = 0.40). **(B)** miR-29c expression was significantly reduced following anti-miR-29a treatment (scr: N = 3; anti-miR-29a: N = 3; unpaired t-test: p < 0.01; Mann–Whitney U test: U = 0, p = 0.10).

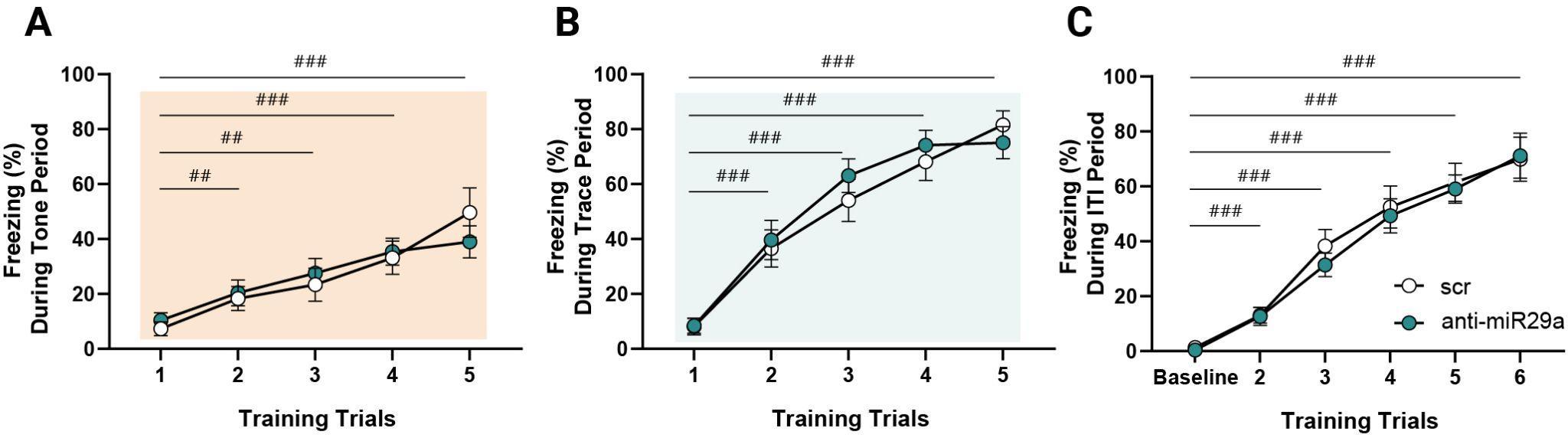

**Supplementary Figure 4. Learning curves during TFC training.** Both anti-miR29a and scr treated mice showed the same learning profile during conditioning for the different TCF training phases. **(A)** Freezing during Tone presentation (Two-way RM ANOVA, main effect of treatment F(1,19) = 8.73x10-4, p = 0.98; main effect of trials F(2.61,49.71) = 23.77, p < 0.0001; trials × treatment interaction F(4,76) = 1.21,p = 0.31). **(B)** Freezing during Trace period (Two-way RM ANOVA , main effect of treatment F(1,19)=0.22, p = 0.63; main effect of trials F(3.43, 65.28) = 58.40, p < 0.0001; trials × treatment interaction F(4,76) = 0.64,p = 0.63). **(C)** Freezing during intertrial interval (ITI, Two-way RM ANOVA, main effect of treatment F(1,19) = 0.14, p = 0.70; main effect of trials F(2.36,44.86) = 78.13, p < 0.0001; trials × treatment interaction F(5, 95) = 0.21, p = 0.95). scr: N=10, anti-miR29a N=11; Tukey’s multiple comparison vs trial 1 or baseline, ##P-value < 0.01, ###P-value < 0.001.

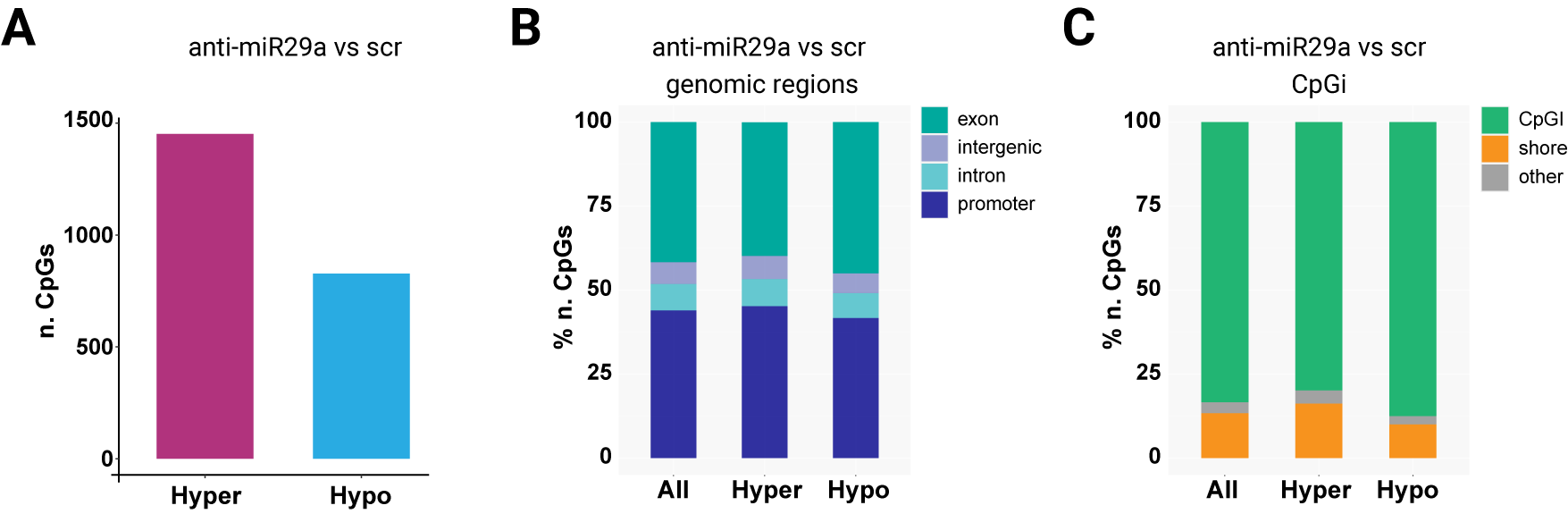

**Supplementary Figure 5. DNA methylation changes in the dorsal hippocampus after miR-29a downregulation in a different cohort of mice. (A)** The number of hyper- and hypo- methylated CpGs. We found 2,279 differentially methylated CpGs (DMCs), with a prevalence of hypermethylation over hypomethylation (1,452 CpGs vs 827 CpGs, p < 10-16, Fisher’s exact test) between anti29a and scr hippocampal treated samples. **(B-C)** DMCs distribution across chromosomes. DMCs exhibited an even distribution across chromosomes and were enriched in CpG island (CpGi).

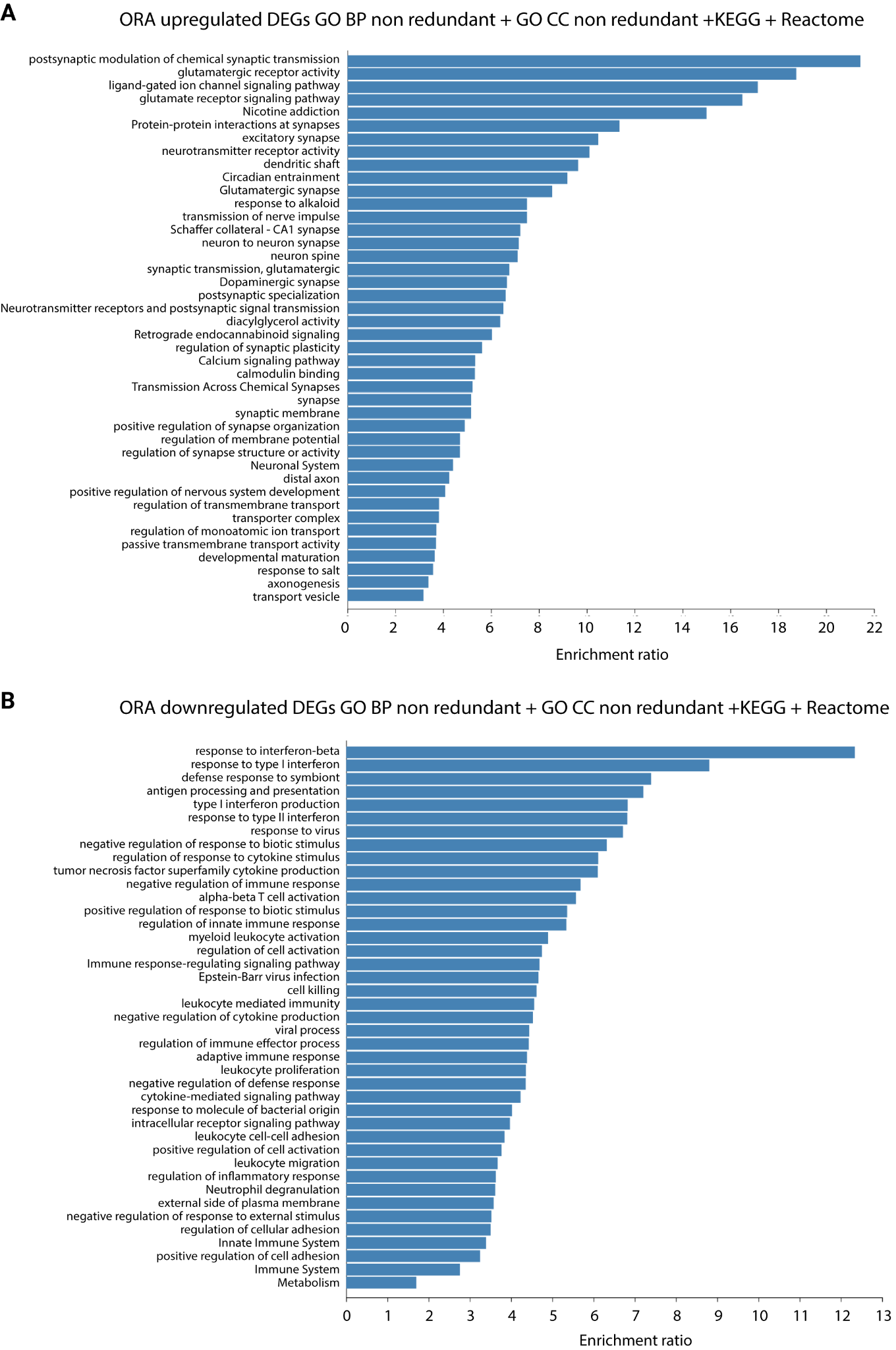

**Supplementary Figure 6.** **Functional enrichment analysis of differentially expressed genes (DEGs).** Over-representation analysis (ORA) was performed on upregulated and downregulated DEGs using non-redundant terms from Gene Ontology Biological Process (GO BP), Gene Ontology Cellular Component (GO CC), KEGG, and Reactome databases. Bar plots show the top enriched categories ranked by enrichment ratio. **(A)** Enriched pathways among upregulated DEGs, predominantly associated with neuronal and synaptic functions, including chemical synaptic transmission, glutamatergic signaling, synapse organization, and neuronal system processes. **(B)** Enriched pathways among downregulated DEGs, mainly related to immune and inflammatory responses, including response to interferon-β, defense response to virus, antigen processing and presentation, cytokine signaling, and leukocyte-mediated immunity.

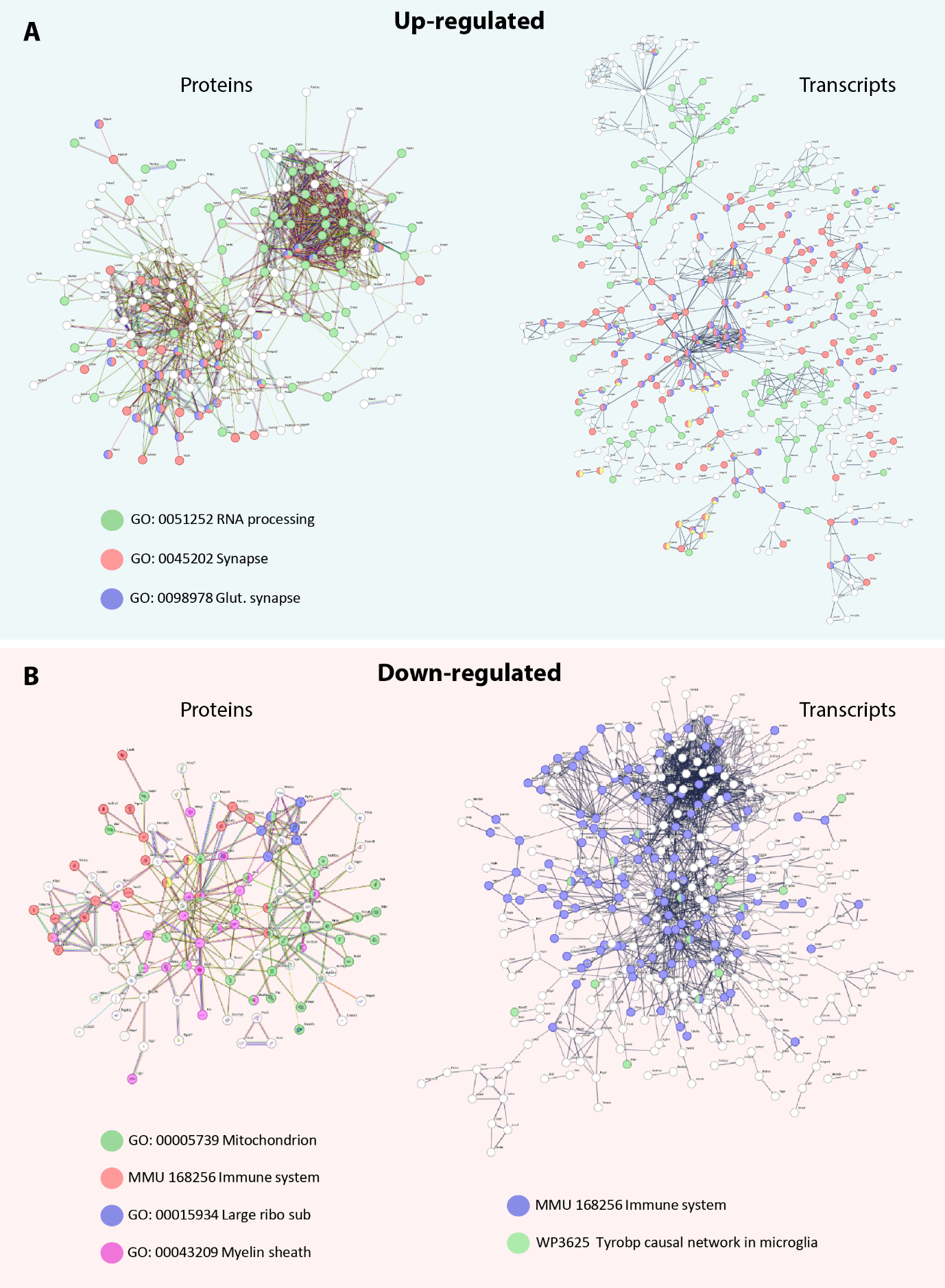

**Supplementary Figure 7. Functional interaction networks of differentially expressed transcripts and proteins following miR-29 downregulation. (A)** Networks of significantly upregulated proteins (left) and transcripts (right). **(B)** Networks of significantly downregulated proteins (left) and transcripts (right). Nodes represent genes or proteins, and edges indicate functional or physical interactions. Colored nodes correspond to genes/proteins annotated in Gene Ontology (GO) or Reactome with the selected terms, codes linking terms to color are indicated in the legends. White nodes represent genes/proteins which are part of the interaction network but are not annotated with any of the highlighted enriched terms.. Network analysis was performed using STRING.

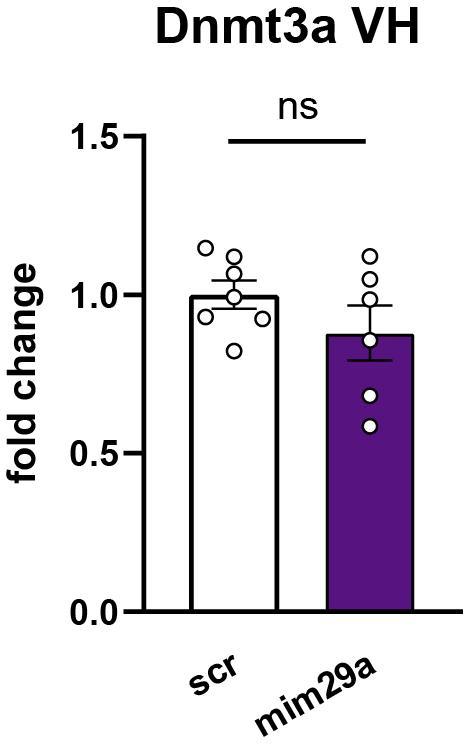

**Supplementary Figure 8.** **Quantification of *Dnmt3a*** **levels in the ventral hippocampus of mim29a and scr treated mice**. **(A)** mimic 29a injection in the dorsal hippocampus did not affect the expression of *Dnmt3a* in the ventral hippocampus confirming the specificity of the injection site within the dorsal hippocampus (Mann-Whitney U test: U =,14 p = 0.37). scr: N = 7, mim29a N = 6. DH = Dorsal Hippocampus; VH = Ventral Hippocampus.

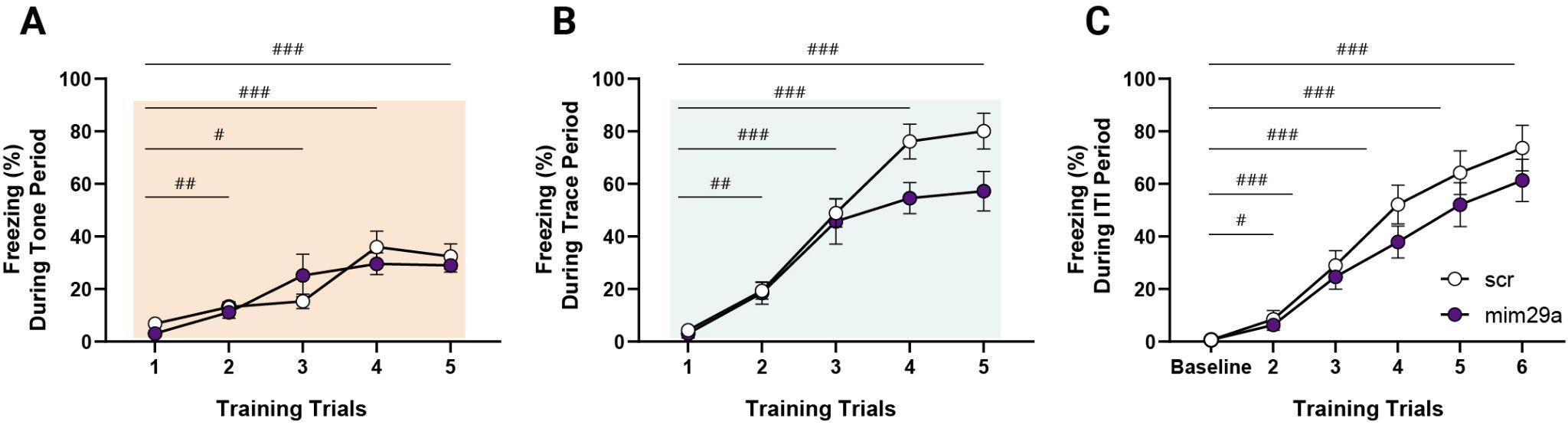

**Supplementary Figure 9. Learning curves during TFC training**. Both mim29a and scr treated mice showed the same learning profile during conditioning for all the different TCF training intervals. **(A)** Freezing during Tone presentation (Two-way RM ANOVA, main effect of treatment F(1,14) = 0.11, p = 0.75; main effect of trials F(2.52,35.24) = 21.48, p < 0.0001; trials × treatment interaction F(4,56) = 1.53, p = 0.20). **(B)** Freezing during Trace period (Two-way RM ANOVA, main effect of treatment F(1,14) = 4.18, p = 0.06; main effect of trials F(3.79,39.14) = 64.76, p < 0.0001; trials × treatment interaction F(4,56) = 2.5, p = 0.053). **(C)** Freezing during intertrial interval (ITI, Two-way RM ANOVA, main effect of treatment F(1,14) = 0.58, p = 0.23; main effect of trials F(2.64,36.99) = 66.44, p < 0.0001; trials × treatment interaction F(5,70) = 0.83, p = 0.53). scr: N = 10, anti-miR29a N = 11; Tukey’s multiple comparison vs. trial 1 or baseline, # P-value < 0.05, ## P-value < 0.01, ### P-value < 0.001.

**Supplementary Table 1: Statistical analysis of sex differences across behavioral and molecular outcomes in anti-miR29a and miR29a mimic experiments.**

LNA-anti-miR29a	miR29a mimic

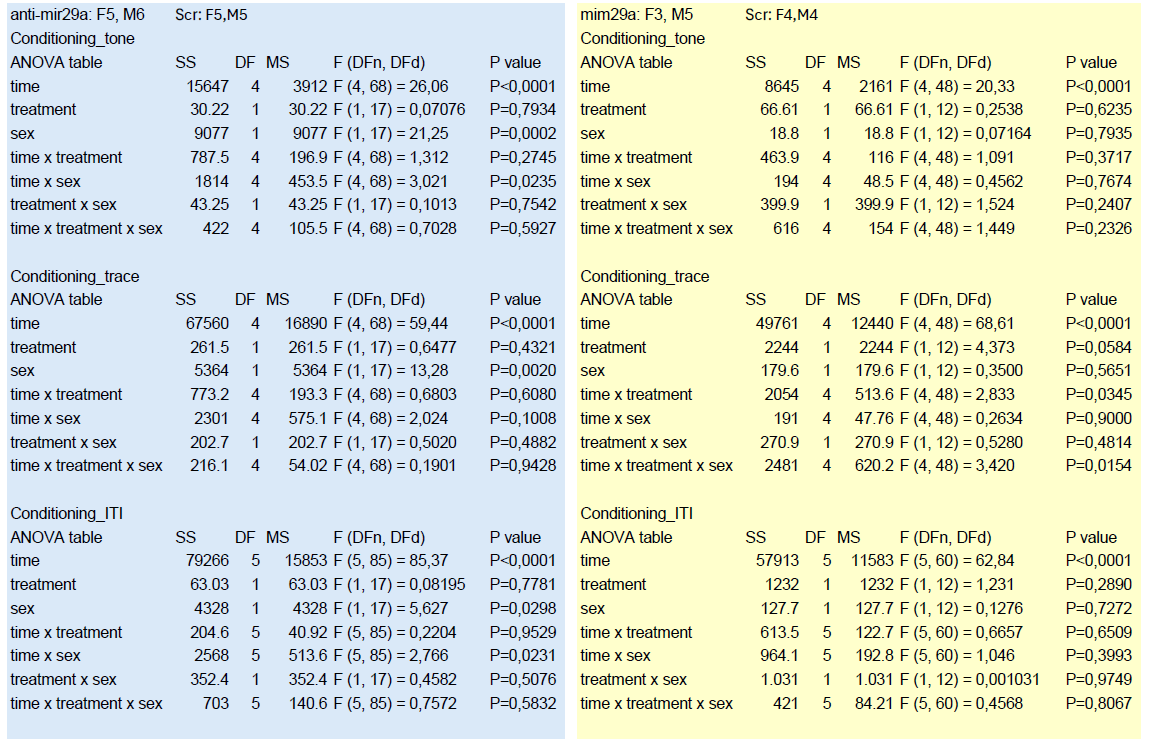

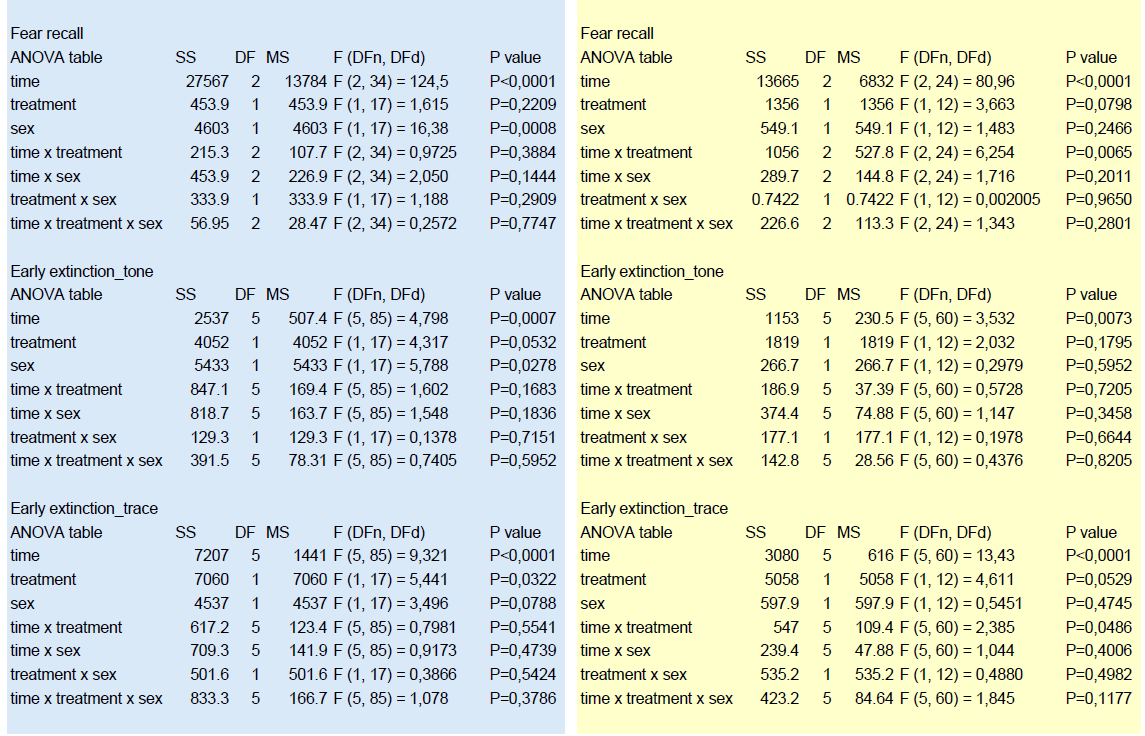

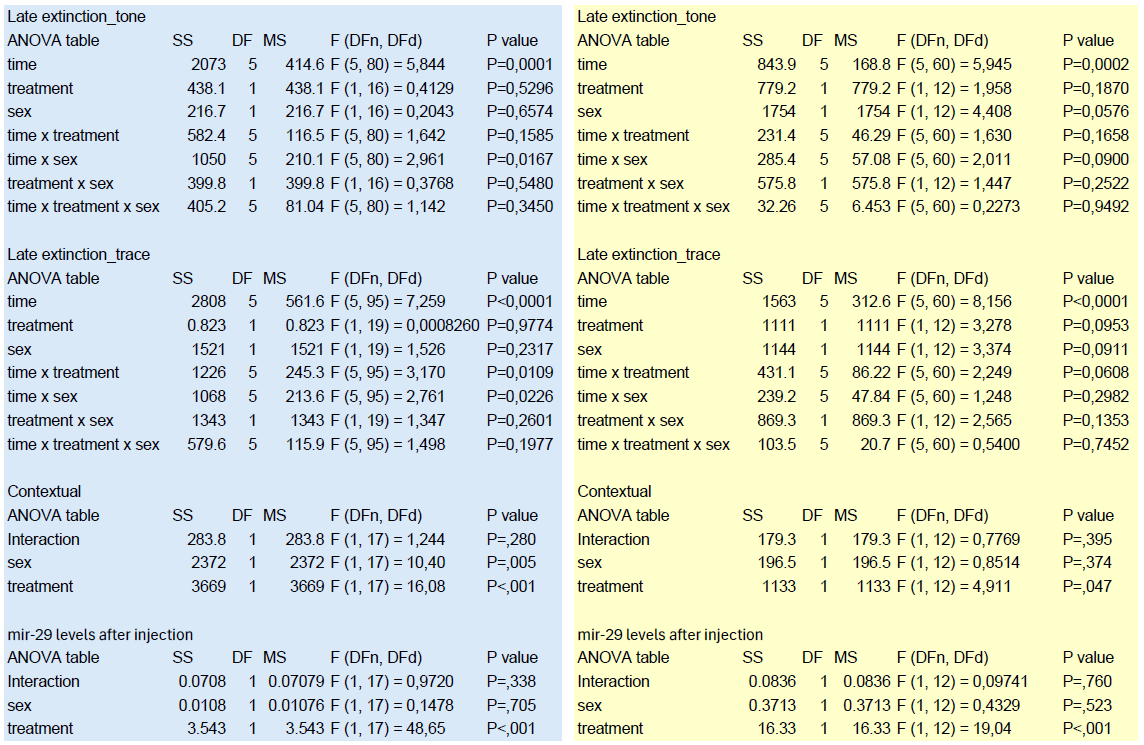

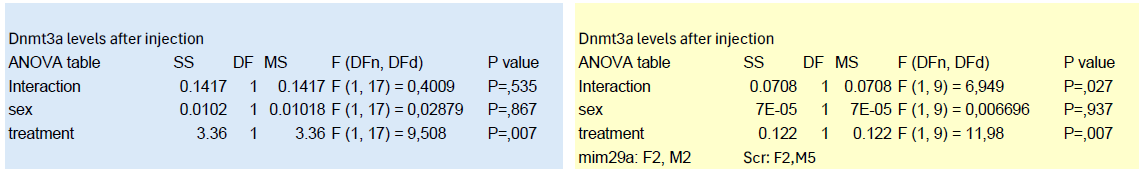
